# Welcome pathogens: transient heat dampens immune responses to acibenzolar-*S*-methyl in apple plants

**DOI:** 10.1101/2025.04.03.647103

**Authors:** Erwan Chavonet, Bao-huynh Nguyen, Christelle Heintz, Raphaël Cournol, Jitka Široká, Ondřej Novák, Romain Larbat, Matthieu Gaucher, Marie-Noëlle Brisset, Florent Pantin, Alexandre Degrave

**Author notes:** Authors for correspondence: Florent Pantin, Alexandre Degrave. Co-first authors (Erwan Chavonet, Bao-huynh Nguyen).

## Abstract

Climate change affects plant-pathogen interactions, with disease outcome varying depending on pathosystem and environmental scenario. In Arabidopsis, a thermo-sensitive module of salicylic acid (SA) signaling makes immunity vulnerable to heat. The potent resistance inducer acibenzolar-*S*-methyl (ASM), an SA analogue that up-regulates transcription of defense genes, could restore plant protection under heat but not core SA signaling. Here, we investigated how high temperature rewires the ASM-induced responses of the apple immune system. We treated apple plants with ASM under contrasting heatwave scenarios and subsequently exposed them to *Erwinia amylovora* (the fire blight bacterium) or *Venturia inaequalis* (the apple scab fungus) while monitoring gene expression. While pre-exposing apple plants to high temperature did not change their susceptibility to pathogens, it drove a loss of ASM-induced protection. Transcriptomic analysis revealed broad dampening of ASM-regulation upon high temperature, for a wide range of biological processes beyond defense. We uncovered thermo-sensitive “resistance” and “susceptibility” marker genes with ASM-responsiveness being critically vulnerable to heat. We concluded that exposure to heatwave prevents ASM from fully mounting its protective responses in apple, not only lowering defenses but also offering more favorable hosting conditions. Our work highlights plant immunity as the joint outcome of resistant and susceptible responses.

**Summary Statement:** We found that heatwaves “disarm” apple’s ability to mount an effective inducible-immunity response against two major diseases, fire blight and apple scab. Heatwaves not only prevent full expression of plant defenses, but also favor a physiological status that is beneficial to the pathogens.

## Introduction

Climate change modifies the many interactions between plants and pathogens. The ‘disease triangle’ framework posits that successful disease establishment requires a virulent pathogen, a susceptible host plant and favorable environmental conditions (Stevens, 1960; Velásquez *et al*., 2018; Roussin-Léveillée *et al*., 2024). Changes in temperature, radiation, relative humidity, soil water content and atmospheric CO_2_ concentration may all affect the physiology and development of plants and pathogens, either directly or through their mutual interaction. For instance, temperature elevation has profound effects on plants, triggering thermomorphogenesis (Quint *et al*., 2016), acceleration of phenology and growth (Parent *et al*., 2010), alteration of carbon balance (Vasseur *et al*., 2011), tissue desiccation (Marchin *et al*., 2022) or even burning (Coupel-Ledru *et al*., 2024), depending on the range and duration of temperature increase, the genetic background and the influence of other environmental factors (Long & Ort, 2010). Temperature elevation may also affect different components of the plant immune system, either positively or negatively (Malamy *et al*., 1992; Zhu *et al*., 2010; Cheng *et al*., 2013; Zhang *et al*., 2017; Richard *et al*., 2020; Qiu *et al*., 2022; Kim *et al*., 2022). In turn, temperature elevation largely affects the growth rate and virulence of microbes, resulting in positive or negative effects depending on the species and strain, with feedback from host responses (Velásquez *et al*., 2018; Roussin-Léveillée *et al*., 2024). Thus, warming may confer more or less chance to a given microbe to meet and infect a susceptible plant, depending on the pathosystem and the environmental scenario (Desaint *et al*., 2021; Chaloner *et al*., 2021).

Focusing on the model pathosystem *Arabidopsis thaliana* / *Pseudomonas syringae*, recent reports have established that plant immunity is vulnerable to heat (28–30°C), through a negative effect on salicylic acid (SA) – one of the major hormones of plant defense (Peng *et al*., 2021). Such vulnerability arises from a thermosensitive pair of master transcription factors (CBP60g and its paralogue SARD1) that fails to induce the expression of SA biosynthesis genes upon heat, making host plants more susceptible (Huot *et al*., 2017; Kim *et al*., 2022; Shields *et al*., 2025). From the pathogen’s perspective, elevated temperature also enhances virulence by accelerating translocation of bacterial effector proteins, leading to more severe disease under heat (Huot *et al*., 2017). However, over- or targeted-expression of *CBP60g* is sufficient to restore thermo-resilient SA production and effective defense (Kim *et al*., 2022), which suggests that loss of host SA synthesis is the major driver of enhanced disease under heat within this pathosystem. Application of acibenzolar-*S*-methyl (ASM, also known as BTH) effectively prevents bacterial infection at warm temperature but, interestingly, does not restore the SA pathway (Huot *et al*., 2017). ASM is a long-standing plant resistance inducer that is converted *in planta* into acibenzolar, a biologically active form that shares functional and structural analogies with SA (Görlach *et al*., 1996; Canet *et al*., 2010a; Tripathi *et al*., 2010). While free or bound SA may accumulate upon ASM application in some species and environmental conditions (e.g. Kuźniak *et al*., 2014; Huot *et al*., 2017; Ullah *et al*., 2019; Ratchaseema *et al*., 2021), there is clear evidence that SA accumulation is dispensable for efficient ASM-induced protection (Friedrich *et al*., 1996; Lawton *et al*., 1996; Canet *et al*., 2010a; Huot *et al*., 2017). By contrast, ASM-induced protection critically requires NPR1 (NONEXPRESSER OF PR GENES 1; Lawton *et al*., 1996; Canet *et al*., 2010a,b; Huot *et al*., 2017), now known as the SA receptor acting as a transcriptional co-activator (Ding *et al*., 2018). It is therefore likely that NPR1 directly recognizes acibenzolar to activate downstream defense genes such as the canonical SA marker *PR1* (*PATHOGENESIS-RELATED GENE 1*). In Arabidopsis at elevated temperature, applying ASM properly initiates NPR1 signaling and yet fails to induce *PR1* expression and feedforward SA accumulation (Huot *et al*., 2017). This suggests that ASM can counterbalance heat-sensitive immunity via other routes than the core SA pathway.

It is noteworthy, yet, that ASM regulates the transcription of hundreds of genes with biological roles beyond defense, for instance by down-regulating genes related to metabolism, photosynthesis or cell division (Huot *et al*., 2017; Warneys *et al*., 2018). Among these processes that are routinely sustained by plants for their physiology and development, some may involve susceptibility factors that are exploited by pathogens to facilitate colonization and propagation *in planta* (Eckardt, 2002; van Schie & Takken, 2014; Gorshkov & Tsers, 2022). Thus, it is perhaps useful to consider plant immunity as the joint outcome of resistant (R) and susceptible (S) responses. In this framework, ASM-induced immunity would result from both the enhancement of defense genes and the repression of susceptibility genes, with different effects on immunity, physiology and development, depending on the species and the environment. To what extent high temperature may affect the different components of ASM-induced immunity remains largely unexplored.

We addressed this question on apple (*Malus domestica*), where ASM effectively controls immunity by eliciting the production of defense compounds such as pathogenesis-related (PR) proteins, and promotes resistance against several pests (Brisset *et al*., 2000; Maxson-Stein *et al*., 2002; Dugé de Bernonville *et al*., 2014; Aćimović *et al*., 2015; Marolleau *et al*., 2017; Warneys *et al*., 2018; Chavonet *et al*., 2022). These include *Erwinia amylovora*, a bacteria responsible for fire blight (Vanneste, 2000), and *Venturia inaequalis*, a fungus responsible for apple scab (Bowen *et al*., 2011). Both pathogens exclusively infect young organs, yet their infection routes and pathogenic mechanisms differ. Following penetration through natural pores or wounds of young leaves or flowers, *E. amylovora* colonizes host apoplast by feeding on intracellular content spilling out from cell death induced by bacterial type III effectors (Zeng *et al*., 2021) and causes progressive necrosis along the stem – potentially leading to tree death. Exudation of ooze droplets from infected tissues is responsible for secondary infections. By contrast, germ tubes from ascospores of *V. inaequalis* penetrate the cuticle of developing leaves or fruits, either directly or through an appressorium, before forming stromata that extract plant nutrients from the subcuticular space. Stromata then produce conidia that are spread by wind and rain, originating secondary infections that occur throughout fruit development, leading to necrotic lesions that render the fruits unmarketable (Bowen *et al*., 2011). Most commercial apple cultivars are susceptible to *E. amylovora* (Kostick *et al*., 2019) and *V. inaequalis* (Gessler *et al*., 2006; Holb, 2007). The application of resistance inducers such as ASM is a promising strategy to manage the protection of susceptible orchards against these two diseases at least. Nonetheless, the efficacy of resistance inducers depends on environmental conditions (Walters *et al*., 2005, 2013; Sandroni *et al*., 2020) – and the components of immunity that are susceptible to the environment remain to be identified.

Here, we found that the efficacy of ASM protection against *E. amylovora* and *V. inaequalis* erodes when apple plants are pre-exposed to transient heat. Such vulnerability to heat was not accompanied by misregulation of master defense modules, contrasting with Arabidopsis. Instead, transcriptomic analyses revealed that transient heat triggered pervasive dampening of ASM-induced transcriptional regulation on a wide range of physiological processes, leaving apple plants less immune. Thus, transient heat ‘imprints’ the physiology of apple plants so that ASM-induced immunity becomes less protective against pathogens in these conditions.

## Materials and Methods

### Plant materials and growth conditions

Experiments were performed on apple seedlings (3–6 leaves) obtained from seeds of open-pollinated trees of cv. Golden Delicious, or on scions (2 to 6-month-old) of Golden Delicious doubled-haploid 13 (GDDH13; Daccord *et al*., 2017) grafted on the MM106 rootstock. Plants were cultivated in semi-controlled greenhouse conditions (natural photoperiod supplemented with artificial light to reach a 16-h photophase, 23–25°C day, 17°C night, watering every 2–4 days) in individual pots filled with a growing medium (Neuhaus Huminsubstrat N4, Klasmann-Deilman France, Bourgoin-Jallieu, France). Both seedlings and grafted plants were chosen for protection experiments because of their overall susceptibility to fire blight and apple scab, and transcriptomic analyses were conducted on GDDH13 grafted scions to take advantage of the high-quality genome sequence of this homozygous genotype (Daccord *et al*., 2017). In all experiments, the youngest unfolded leaf at time of treatment application was identified by attaching a small clip to the petiole just before spraying ASM or water.

### Temperature and chemical treatments

Temperature scenarios were applied in environmental chambers FitoClima 600 (ARALAB, Rio de Mouro, Portugal) with controlled temperature, relative humidity (RH) and radiation provided by white, blue and red LED tubes. In a first set of experiments, plants were exposed to constant temperature (20°C) and RH (60%) day and night (12-h photoperiod at 300 µmol m^−2^ s^−1^). Heat was imposed by switching temperature to 35°C during 24 h or during the 12-h photophase, depending on the experiment as described in the Results section. In the following experiments, the environmental scenarios were refined to match natural fluctuations, based on the average temperature dynamics on the 11 hottest days (peak at 34–36°C) between March and July 2016–2018 in an experimental orchard (UE449 HORTI, INRAE, Beaucouzé, France) adjacent to our research center (Fig. S1). A typical heat day was designed through piecewise linear approximation of temperature, relative humidity and irradiance over time, with a 4-h heat plateau in the afternoon (“Short” settings”) to be compared to standard conditions for apple tree growth in the same French area (“Control” settings”) and to an artificially extended 12-h heatwave (“Long” settings) (Fig. S2). Nighttime temperature was set at 18°C until 8:00 am, followed by a 2-h ramp and a 12-h plateau at 21°C and 35°C for the “Control” and “Long” settings, respectively, or a 6-h ramp and a 4-h plateau at 35°C for the “Short” setting. Finally, a 2-h (“Control” and “Long”) or a 6-h (“Short”) decreasing ramp was applied to reach 18°C at midnight. To match natural variations, RH was covaried with temperature, reaching 80%, 60% and 40% when temperature was 18°C, 21°C and 35°C, respectively (Fig. S2). The same daily fluctuations in radiation were applied to the three regimes, based on the radiation measured in the orchard. Lighting progressively started at 8:00 am to reach 300 µmol m^−2^ s^−1^ between midday and 4:00 pm, and progressively declined to switch off at 8:00 pm. The number of days of heat vs. control temperatures depended on the objective of experiment, and each scenario is described in Results for each experiment.

Acibenzolar-*S*-methyl (ASM) solutions were prepared from the commercial product Bion® 50WG (Syngenta, Basel, Switzerland; 50% of ASM) in reverse osmosis water at 0.4 g L^−1^. The same water was systematically used as control. Aerial parts of plants were sprayed until incipient runoff with a spray gun Aeryo-1.4 (Deltalyo, Mably, France) between 8:00 am and 10:00 am on the day selected for the treatment. The timing of treatment in relation to temperature scenario depended on the experiment and is described in Results.

### Quantification of ASM in treated leaves

The morning of the experiment, seedlings from the greenhouse were sprayed with ASM (t_0_) and immediately transferred in ARALAB chambers at 20°C or 35°C (60% RH, 300 µmol m^−2^ s^−1^ PAR). After 6 h, when all leaves were unequivocally dry, the youngest unfolded leaf was harvested on 2 plants, rinsed 3 times in water, dried with paper towel, pooled in an Eppendorf and frozen in liquid nitrogen. This protocol was repeated on the leaf insertion rank immediately below. For each leaf rank and each temperature, 10 replicates were obtained. Pooled samples were crushed to a fine powder with mortar and pestle. Fresh leaves powders (50 mg) were suspended in 500 µL of methanol spiked with 0.2 µg mL^−1^ of [^2^H_6_]ABA as internal standard. After 10 s vortexing, mixtures were stored for 15 h at −20°C and then centrifuged at 10000 g for 10 min at 4°C. The supernatant was collected and the pellet extracted a second time with 500 µL of methanol. The two combined supernatants were then evaporated using a vacuum concentrator. Dried extracts were resuspended in 200 µl of methanol, filtered (0.2 µm) and transferred to vials before LC-MS analysis.

Chromatographic analyses were performed on a Vanquish Flex UHPLC system equipped with a quaternary solvent manager, an autosampler and a temperature-controlled column. ASM and metabolites contained in the extracts (1 µL) were separated on a HSS-T3 column (100 × 2.1 mm, 1.8 µm) (Waters, Milford, MA, USA) using a gradient of mobile phase composed of water + 0.1% formic acid (A) and methanol + 0.1% formic acid (B) at a flow rate of 300 μL min^−1^. The samples were kept at 10°C and the column was maintained at 25°C. The elution program consisted in a first 2-min step with 0% B, then a linear ramp from 0% to 100% B in 18 min, followed by a 3-min step with 100% B, before re-equilibrating the column in the initial conditions for 7 min. The samples were analyzed randomly.

MS^1^ detection was performed on a TSQ Quantis Plus^TM^ (ThermoFisher Scientific, Bremen, Germany) mass spectrometer in positive and negative electrospray ionization (ESI) modes. The capillary voltages were set at 3 kV and 2.75 kV for positive and negative modes, respectively. The source gases were set (in arbitrary unit min^−1^) to 40 (sheath gas), 10 (auxiliary gas) and 0 (sweep gas) and the vaporizer temperature was 280°C. Full scan MS^1^ spectra were acquired from 50 to 700 *m/z* with a scan rate of 1000 Da s^−1^. ASM was quantified in selective ion monitoring (SIM) mode by following ion at *m/z* 211 in positive mode. Acibenzolar acid was monitored at *m/z* 179 in negative mode together with the internal standard [^2^H_6_]ABA at *m/z* 269. ASM, acibenzolar acid and [^2^H_6_]ABA pure standards eluted at 18.1 min, 14.1 min and 19.7 min, respectively, with the above-mentioned conditions. ASM and acibenzolar acid quantities in dry laminas were determined using respective standard curves. Control leaves were also harvested at t_0_ (without any spray) from independent plants in the greenhouse, and all samples were below the detection limit.

### Physiological measurements

The morning of the experiment, each plant was first characterized in the greenhouse. Stem height (from the cotyledon to the apical meristem) and lamina length (youngest unfolded leaf, just identified using a clip) were measured using a ruler. Stomatal conductance (g_s_) of the clipped leaf and of the leaf immediately below (more mature and less susceptible to *E. amylovora* and *V. inaequalis*) was determined using a porometer (LI-600; Li-COR Inc., Lincoln, NE, USA). Plants were then treated with ASM or water (t_0_) and immediately placed in ARALAB chambers at 12-h photoperiod, 250–300 µmol m^−2^ s^−1^ PAR, and 60% RH. Air temperature was maintained at either 20°C or 35°C for 24 h. g_s_ of both leaves were measured again at 24, 48 and 72 h after treatment. Stem height and lamina length of the clipped leaf were measured again at 72 h after treatment and absolute elongation rates were computed as the difference between initial and final lengths divided by the time interval.

### Pathogen strains and protection experiments

Plants were inoculated by the CFBP1430 strain of *E. amylovora* (Paulin & Sampson, 1973). Inoculum was suspended at 10^8^ cfu mL^-1^ in reverse osmosis water from an exponential growth phase culture cultivated overnight at 26°C on solid King’s B medium (King *et al*., 1954). Inoculation was carried out in greenhouse conditions (1 or 3 d following ASM treatment for seedlings, and 3 d after ASM treatment for grafted plants), by cutting two-thirds of the two youngest unfolded leaves (perpendicularly to the midrib) with scissors previously dipped in the inoculum. Progressions of necroses from the inoculated leaves to the stems, through the midrib and petiole, were assessed 21 or 28 d post-inoculation (dpi) for seedlings or grafted plants, respectively. Disease incidence was calculated as the percentage of diseased plants, considering that a plant was diseased when typical fire blight necrosis reached the stem, whereas it was considered healthy if no necrosis occurred or if necrosis was restricted to the inoculated leaf (including its petiole) at the end of the experiment.

The *V. inaequalis* strain 104 (Guillaumès *et al*., 1995) was prepared in reverse-osmosis water from scabbed apple leaves previously stored at −80 °C. The suspension was roughly filtered to remove debris, adjusted to a concentration of 1.5 × 10^4^ conidia mL^-1^ and sprayed onto apple seedlings until incipient runoff. To facilitate conidial germination and fungus penetration, plants were then placed in a controlled environment room in the dark, at 18°C and with humidity maintained above 90% for 48 h. Light was then switched on (16-h photoperiod), and humidity reduced to 70% while temperature was maintained at 18°C. Symptoms were assessed 21 d after inoculation on the youngest unfolded leaf at the time of ASM application. The proportion of leaf area affected by sporulating lesions was scored according to a 7-category ordinal scale (Calenge *et al*., 2004). Disease incidence was calculated as the percentage of diseased plants, considering that a plant was diseased when visible sporulation occurred on the target leaf (category > 0).

The numbers of independent trials and biological replicates are indicated in the figure legends.

### Sampling for gene expression and protein analyses

Sampling for gene expression or protein analysis was carried out by pooling the youngest unfolded leaf (or its distal two-thirds cut out while inoculating *E. amylovora*) from 1 to 5 plants to form one replicate. Samples were immediately frozen in liquid nitrogen and stored at -80°C before processing. For the retrospective transcriptional analysis, leaves were individually sampled while inoculating with *E. amylovora*, and pooled *a posteriori* depending on the healthy or diseased fate of their corresponding plant. This was done through hierarchical clustering analysis based on symptom severity (necrosis length) scored at 7, 14 and 21 d after inoculation, where the most distant groups were selected to distinguish between healthy and diseased plants. The genetic material (Golden Delicious seedling or GDDH13 grafted plants) as well as the number of biological replicates and independent experiments are indicated in the figure legends.

### Microarray procedure

Leaves were ground in fine powder (Retsch, Haan, Germany) and total RNA was extracted from about 200 mg using the Nucleospin RNA Plant Kit (Macherey-Nagel, Düren, Germany) according to the manufacturer’s protocol. Samples purity and concentration were assayed with the Agilent 2100 bioanalyzer system and the RNA 6000 Nano LabChip kit (Agilent Technologies, Waldbroon, Germany) according to the manufacturer’s recommendation. Two hundred ng of RNA were amplified and labelled as described in Perrin *et al*. (2020). Labelled cDNA were prepared for hybridization according to the manufacturer’s procedure with the Gene Expression Hybridization Kit (Agilent, Waldbronn, Germany) and the Hi-RPM gene Expression Hybridization Kits (Agilent, Waldbronn, Germany) and hybridized on AryANE v1.1 slides (Agilent, design n°85275) in incubation chambers for 17 h at 65°C. After washing, slides were scanned at 3 µm with an Innoscan 710 using Innoscan v3.4.7 software (Innopsys, Carbonne, France).

### Transcriptional marker analysis

Leaves were ground and total RNA were extracted as described above in the microarray procedure, except that RNA purity and concentration were assessed with a NanoDrop One (Thermo Fisher Scientific, Waltham, MA, USA). Reverse transcription (RT) and quantitative polymerase chain reaction (qPCR) were performed as described in Chavonet *et al*. (2022). The primers and their targets designed for this study are listed in Table S2; while the remaining ones are described in Dugé de Bernonville *et al*. (2014). The relative changes in gene expression were calculated using the 2^−ΔΔ^*^C^*^T^ method (Schmittgen & Livak, 2008) with normalization against three internal reference genes (Vandesompele *et al*., 2002). The mean expression value calculated from samples obtained on water-treated plants exposed to control temperature (or 3-d heatwave for the experiments on heat wave duration and positioning) was used as the calibrator to compute the relative expression as log_2_ ratios.

### Protein extraction, separation and immunodetection

Two hundred mg of deep-frozen leaf powder were used for phenolic extraction of protein and their subsequent solubilization, as described in Chavonet *et al*. (2022). Ten µg of total protein extracts were separated on 1D SDS-PAGE, electroblotted onto 0.45 mm polyvinylidene difluoride (PVDF) membranes (immobilon-P, Milli-pore Corp., Bedford, MA, United States) as described in Warneys *et al*. (2018). Immunodetection of target proteins was performed using the rabbit primary antibodies described in Chavonet *et al*. (2022) using the procedure from Warneys *et al*. (2018).

### Quantification of salicylates

Salicylic acid (SA), SA glucoside (SAG) and SA glucose ester (SGE) in laminas of apple seedlings were quantified in precisely weighed freeze-dried samples (1.8 ± 0.02 mg DW). Samples were extracted in 1 mL of 1 mol L^−1^ formic acid in 10% aqueous methanol with a stable isotope-labeled internal standard ([^2^H_4_]SA at 20 pmol per sample), following Široká *et al*. (2022). Extracts were purified with HLB Oasis® SPE columns (Waters) as described by Floková *et al*. (2014). SA was analysed on an Agilent 1290 Infinity LC system coupled to a 6490 Triple Quadrupole LC/MS, following Široká *et al*. (2022). SAG and SGE were determined from the same samples on an ACQUITY UPLC H-Class PLUS system linked to a Xevo TQ-S micro triple-quadrupole MS (Waters), operating in negative ionization multiple reaction monitoring mode (quantification: 299 > 137, collision energy 12 eV, cone voltage 30 V; confirmation: 299 > 93, collision energy 34 eV, cone voltage 28 V). Separation was performed on an Acquity UPLC BEH Shield C18 column (2.1 × 150 mm, 1.7 µm, Waters) at 40°C. The mobile phase consisted of 0.3% aqueous formic acid (A) and methanol (B), with a flow rate of 0.3 mL min^−1^ and a linear gradient from 5% to 98% B within 10 min. The determination of SAG and SGE was semi-quantitative, using [^2^H_4_]SA as internal standard.

### Data analysis

Data analysis was performed using R (version 4.2.0; R Development Core Team, 2025) and RStudio (version 2024.09.0; R Studio Team, 2025) software except for Clustering Affinity Search Technique (CAST). Multiple comparisons on proportion of diseased and healthy plants were performed using pairwise Fisher’s exact tests of the “rstatix” package (version 0.7.2; Kassambara, 2023a). Two-way ANOVA and estimation of the effect sizes (partial eta squared, η^2^_p_) of individual factors (treatment and temperature) and their interaction were performed using the same package. Multiple comparisons on expression values were performed with a Tukey’s HSD test from package “stats” (R Development Core Team, 2025). Multiple comparisons on growth and conductance data were performed with pairwise t-tests adjusted by the Bonferroni method using “rstatix” (Kassambara, 2023a). Statistical differences for multiple comparisons were established with a risk threshold of 5% designated by different letters using package “rcompanion” (version 2.4.36; Mangiafico, 2024). For microarrays, data normalization and statistical analyses were performed using the LIMMA 3.10.1 package (Smyth, 2005) from the Bioconductor project, following Celton *et al*. (2014). Briefly, after extraction of the scan values with Mapix (version 8.2.5; Innopsys, Carbonne, France), the data were normalized with the lowess method, and differential expression analyses were performed using the lmFit function and Bayes moderated t test. Genes were considered differentially expressed (DEGs) for |log2 ratios| ≥ 1 and p-value ≤ 0.05. Pathway enrichment analysis was performed on all genes dataset without applying filtering with the Gene Set Enrichment Analysis (GSEA, 10000 permutations) by ranked genes by their log_2_ fold-change ratio (Subramanian *et al*., 2005) with Benjamini and Hochberg procedure for controlling false discovery rate (Hung *et al*., 2012), available with the package “ClusterProfiler” (version 4.4.4; Wu *et al*., 2021). CAST was performed on genes corresponding to DEGs in at least one of the comparisons by the software MeV (Multi Experiment Viewer) with Pearson correlation distance metric and an affinity threshold of 0.95 (Ben-Dor *et al*., 1999; Saeed *et al*., 2003). Graphical representations were generated using the package “ggplot2” (version 3.5.1; Wickham, 2016) associated with “ggpubr” (version 0.6.0; Kassambara, 2023b). Heatmaps were performed with “ComplexHeatmap” (version 2.18.0; Gu, 2022). Principal Component Analysis (PCA) were carried out with packages “FactoMineR” (version 2.8; Lê *et al*., 2008) and “factoextra” (version 1.0.7; Kassambara & Mundt, 2020).

## Results

### High temperature impedes the protecting effect of ASM against fire blight and apple scab

To explore the interaction between high temperature and ASM-induced immunity by excluding any direct effect of temperature on pathogen growth or virulence, we targeted ASM and/or heat applications prior to pathogen inoculation. We first sprayed apple seedlings with ASM (or water) and immediately exposed them to a sudden temperature elevation at 35°C during 24 h (or to a 20°C constant regime) as depicted in Fig. 1a (left part). Transient heat and ASM treatments had no obvious impact on plant phenotype, as illustrated by growth or stomatal conductance that remained unaffected (Fig. S3), either on the youngest unfolded leaf at the time of ASM application (rank N) or in the leaf immediately below (rank N-1). Three days after this dual prior treatment, when leaf N had largely developed ontogenetic resistance, we inoculated the newly unfolded leaf (rank N+1) with *E. amylovora* before assessing fire blight symptoms for three weeks. In control conditions (water, 20°C), 82% of plants showed stem necrosis after 21 dpi (Fig. 1b). In the ASM treatment, this proportion dropped to 50%. The variability in basal- and ASM-induced-resistance, consistent with previous studies (Brisset *et al*., 2000; Dugé de Bernonville *et al*., 2014; Chavonet *et al*., 2022), could arise from inherent genetic variability in the seedling population, gradients in the microclimate, and/or subtle differences in the developmental stage of the inoculated leaf. High temperature *per se* did not enhance the susceptibility of apple plants to fire blight in water-treated seedlings (Fig. 1b). However, high temperature abolished the protecting effect of ASM that was observed at control temperature, with 68% of plants showing stem necrosis. Similar results were obtained when seedlings were exposed to a close environmental scenario and further inoculated with *V. inaequalis* (Fig. 1c).

**Fig. 1.**
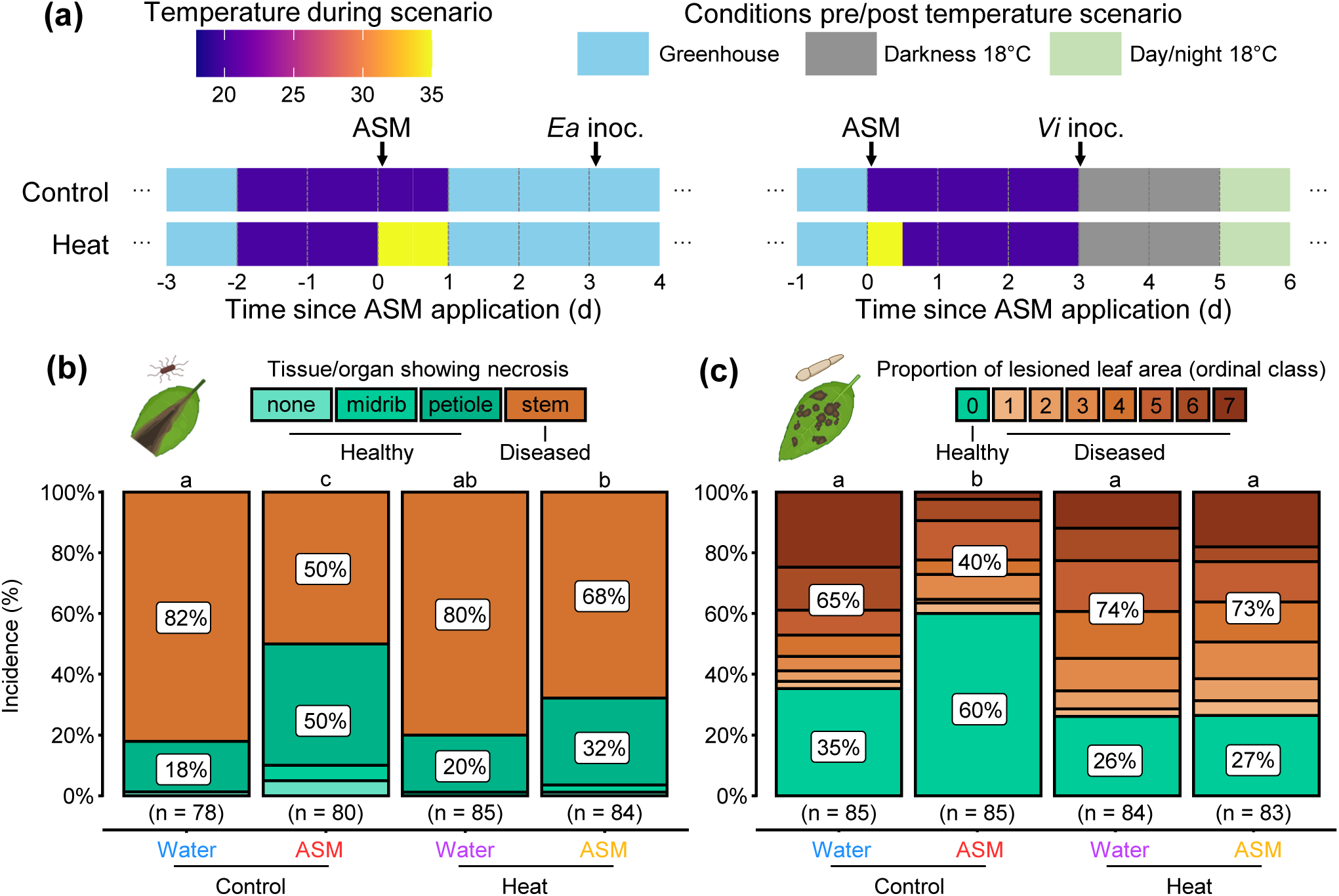
Protection provided by ASM to apple seedlings against *E. amylovora* and *V. inaequalis* is impaired by transient exposure to heat. **(a)** Schematic representation of the experimental protocol applied to plants for *E. amylovora* (left) and *V. inaequalis* (right) disease assessment (details in Methods). Air temperature during the contrasting thermal scenario (period in the growth chamber) is color-coded according to the color map. Arrows indicate the application of ASM (or water) treatment and pathogen inoculation on the youngest unfolded leaf. **(b)** Stacked bar chart showing the proportion of healthy (asymptomatic on stem) and diseased (symptomatic on stem) seedlings inoculated with *E. amylovora*, 21 d post-inoculation. **(c)** Stacked bar chart showing the proportion of healthy (no visible sporulation) and diseased (visible sporulation) seedlings inoculated with *V. inaequalis*, 21 d post-inoculation. Data represent 78 to 85 biological replicates from 2 independent trials for each pathogen (n = total number of plants). Statistical significance was determined using pairwise Fisher’s exact tests; same letters indicate groups that are not significantly different. Icons created in BioRender.com.

To check if this loss of ASM efficiency at high temperature could be observed in more realistic environmental conditions, we devised new scenarios based on records from a local meteorological station that corresponded to heatwave periods (Methods). We varied the duration of heat (numbers of days in the heatwave, and number of heat hours within a day) and inoculated the youngest unfolded leaf with *E. amylovora* systematically 1 d after ASM treatment (Fig. 2a). We found that applying a short heatwave with a 4-h plateau at 35°C during the photophase for 3 d induced a large loss of ASM-efficiency, similar to that in the previous experiment (Fig. 2b). Applying the same heatwave for only one day before inoculation was not sufficient to impede the effect of ASM. Nonetheless, when the heat plateau was extended to 12 h for a single day, ASM lost again most of its protecting effect. Altogether, these results show that accumulation of high temperatures prior to inoculation impairs the ability of apple plants to mount an effective ASM-induced protection against *E. amylovora* and *V. inaequalis*.

**Fig. 2.**
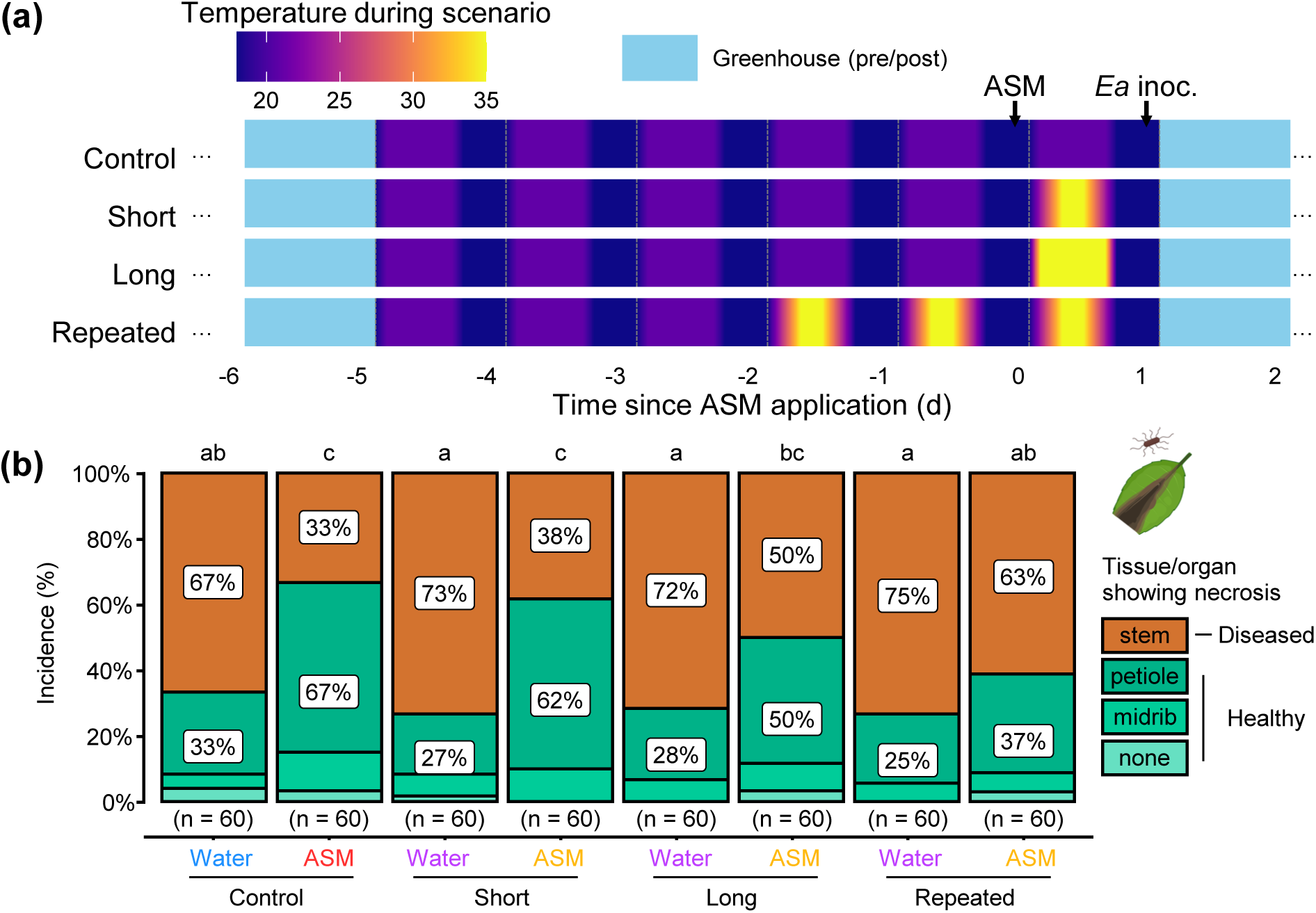
Accumulation of high temperatures prior to inoculation impairs the ability of apple seedlings to mount an effective ASM-induced protection against *E. amylovora*. **(a)** Schematic representation of the experimental protocol (details in Methods). Air temperature during the contrasting thermal scenario (period in the growth chamber) is color-coded according to the color map. Arrows indicate the application of ASM (or water) treatment and pathogen inoculation on the youngest unfolded leaf. **(b)** Stacked bar chart of seedlings showing the proportion of healthy (asymptomatic on stem) and diseased (symptomatic on stem) seedlings inoculated with *E. amylovora*, 21 d post-inoculation. Data represent 60 biological replicates from 3 independent trials (n = total number of plants). Statistical significance was determined using pairwise Fisher’s exact tests; same letters indicate groups that are not significantly different. Icons created in BioRender.com.

### High temperature does not impair ASM uptake or SA metabolism

One trivial explanation might be that high temperature prevents correct uptake of ASM, due to e.g. enhanced evaporation and thus reduced leaf humectation. However, ASM content in treated leaves of apple seedlings 6 h after spraying (after complete evaporation) was similar at 20°C and 35°C (Fig. 3a). The quantity of ASM absorbed per leaf mass was similar between the youngest unfolded leaf (rank N) and the leaf immediately below (rank N-1), showing homogenous and efficient spray. Moreover, acibenzolar acid, the first catabolite of ASM that is biologically active (Tripathi *et al*., 2010), was significantly enhanced at 35°C in both leaf ranks (Fig. 3b). This shows that ASM uptake and metabolism are operational at high temperature.

**Fig. 3.**
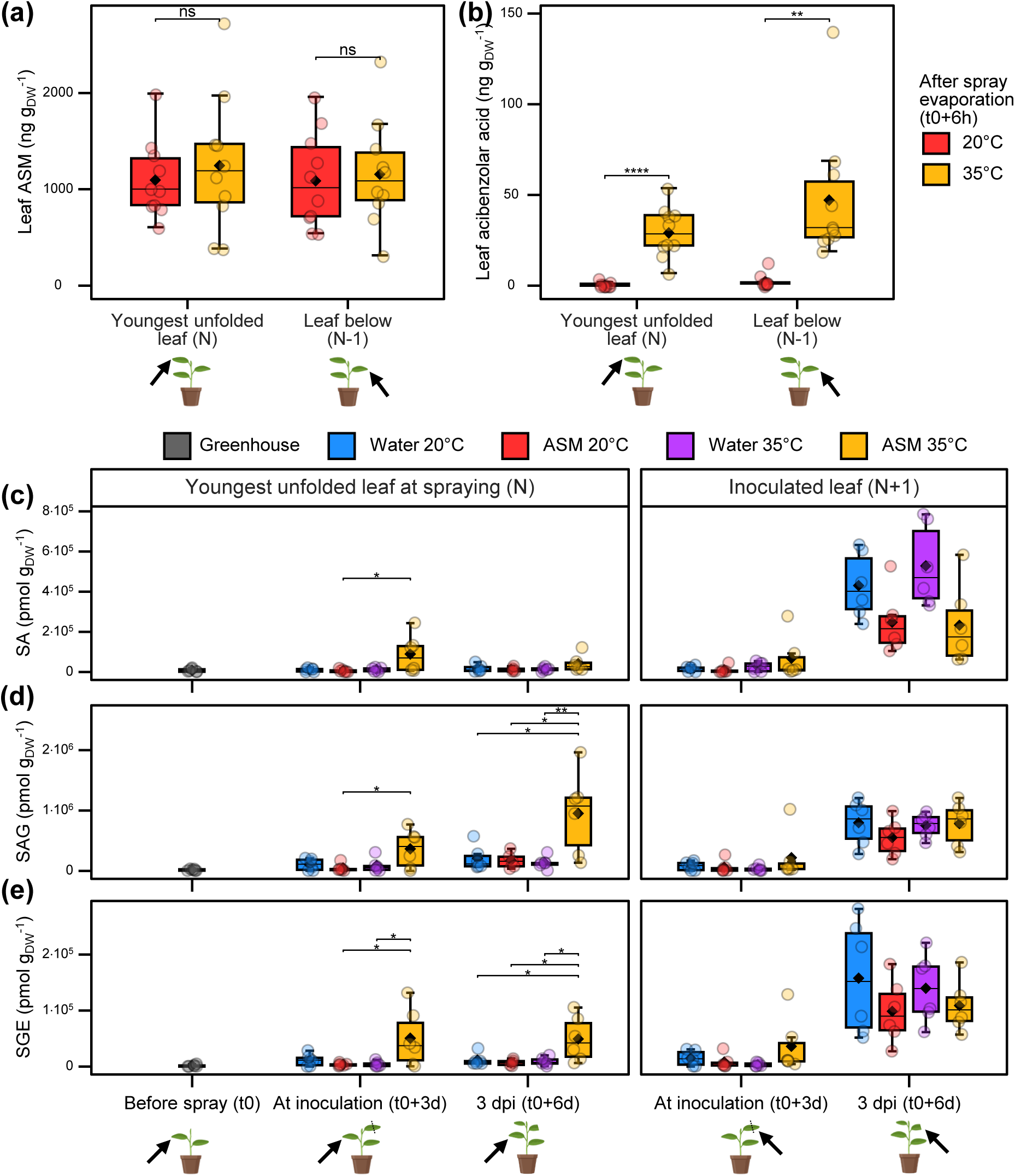
High temperature does not alter ASM uptake or SA metabolism in apple seedlings. **(a)** and **(b)** Content of ASM and acibenzolar acid (biologically-active catabolite) in the lamina of ASM-treated leaves (n = 10, each pooled from 2 different plants). Apple seedlings were sprayed with ASM (t_0_) and immediately transferred into a chamber at 20°C or 35°C. The youngest fully unfolded leaf (rank N) and the leaf below (rank N-1) were sampled after 6 h (when the sprayed solution had evaporated) and thoroughly rinsed before quantification of **(a)** ASM and **(b)** acibenzolar acid. **(c)**, **(d)** and **(e)** Content of salicylates in the lamina of water- or ASM-treated leaves (n = 6, each pooled from 8 different plants). Apple seedlings were sprayed with water or ASM (t_0_) and immediately transferred into a chamber at 20°C or 35°C for 24 h, before being placed in a greenhouse. Three days after spraying (t_0_+3d), half of the plants was sampled for the youngest fully unfolded leaf at time of spraying (rank N) and the leaf above (rank N+1), while the other half was inoculated with *E. amylovora* at leaf N+1. At three days post-inoculation (3 dpi, t_0_+6d), the inoculated leaf N+1 and leaf N were harvested. A batch of plants was also harvested for leaf N just before t_0_, in the greenhouse where the seedlings were grown. The samples were quantified for **(c)** salicylic acid (SA), **(d)** SA glucoside (SAG) and **(e)** SA glucose ester (SGE). ns: not significant, *: P<0.05, **: P<0.01, ***: P<0.001, ****: P<0.0001.

We also checked whether transient heat could alter SA metabolism, as observed in Arabidopsis (Huot *et al*., 2017; Kim *et al*., 2022). In apple seedlings exposed to control temperature, ASM did not induce SA accumulation 3 d after application, either in the youngest unfolded leaf at time of ASM spraying (Fig. 3c, rank N, left), or in the leaf above that had unfolded and was then inoculated (Fig. 3c, rank N+1, right). After 3 d again, the inoculated leaf N+1 showed a large SA accumulation in both water- and ASM-treated plants, consistent with a progression of the bacteria eliciting defense within the leaf tissues, while the proximal, asymptomatic leaf N maintained a basal SA level (Fig. 3c). Transient (24-h) exposure to heat did not affect SA content in water-treated plants the day of inoculation, and ASM-treated plants tended to show enhanced SA, yet a large variability was observed. Three days after inoculation, SA content in either leaf was not affected by the prior temperature treatment. The conjugates SA glucoside (SAG) and SA glucose ester (SGE) essentially showed the same variations as SA (Fig. 3d and e). Thus, there was no apparent link between leaf SA content and ASM-induced protection.

### High temperature does not prevent ASM-induction of defense markers

Our results on apple plants contrast with the Arabidopsis / *Pseudomonas syringae* model pathosystem, where high temperature enhances plant susceptibility to the bacteria but does not affect ASM-induced protection (Huot *et al*., 2017). Nonetheless, in Arabidopsis again, exposure to high temperature for 1 d before and after ASM application inhibits ASM-induced expression of defense genes, including *PR1* (Huot *et al*., 2017). Such inhibition was also observed in rapeseed, tomato and rice (Kim *et al*., 2022). Thus, high temperature might prevent the transcriptional induction of defense genes by ASM in apple plants, potentially underlying the loss of ASM-induced protection. To check this hypothesis, we measured the expression of 29 marker genes covering a broad range of defense mechanisms, including *PR1* (Dugé de Bernonville *et al*., 2014; Chavonet *et al*., 2022), on apple seedlings sprayed with ASM or water and exposed to transient heat or not. Three days after ASM/water treatment, we sampled individual young leaves while inoculating with *E. amylovora*, and kept track of whether the corresponding plants remained healthy or turned diseased. Surprisingly, ASM efficiently induced the transcription of defense markers whatever the temperature scenario, some of the genes being even more induced by ASM at high temperature than at control temperature (Fig. 4a). Thus, the transcriptional induction of defense marker genes by ASM was uncoupled from plant protection at high temperature. Moreover, the expression profile of defense markers at time of inoculation did not account for posterior disease symptoms, as no major difference in gene transcription was observed in plants with healthy or diseased fate (Fig. 4a). This suggests that transcriptional up-regulation of defense genes at the infection site is necessary but not sufficient for effective immunity.

**Fig. 4.**
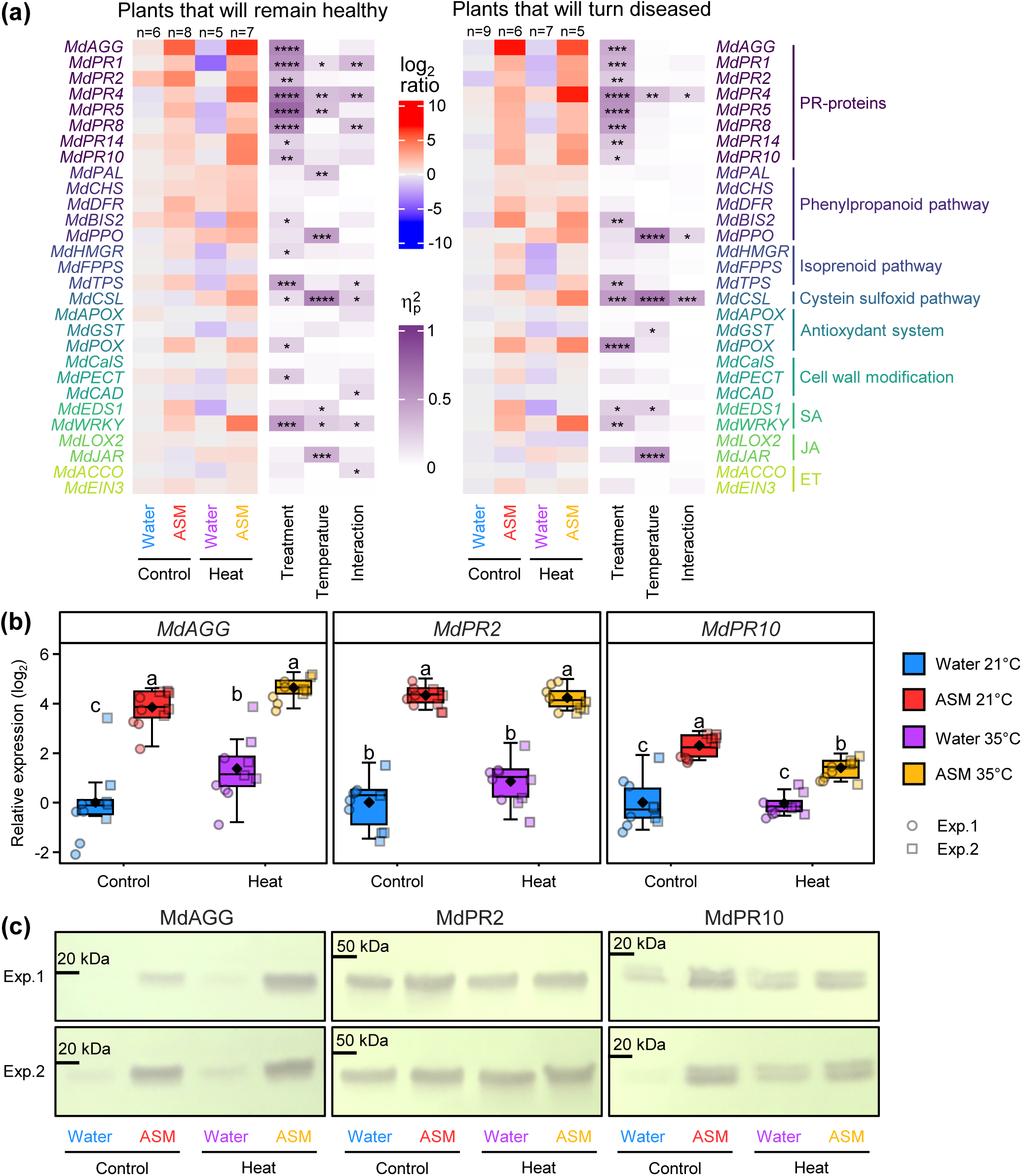
High temperature does not prevent ASM-induction of defense markers. **(a)** Retrospective transcriptional analysis of 29 marker genes covering a broad range of apple defense mechanisms. Following a scenario similar as in Fig. 1a, young leaves of apple seedlings were individually sampled while inoculating with *E. amylovora*, and grouped *a posteriori* depending on the healthy or diseased fate of their corresponding plant. Heatmaps show relative gene expression and ANOVA effect sizes (η^2^_p_) of treatment, temperature and their interaction, as well as their respective significance level (*P<0.05, **P<0.01, ***P<0.001, ****P<0.0001), for both types of plants. Expression values of the average of water-treated leaves from healthy and diseased plants exposed to 20°C were used for the calculation of the log_2_ ratio. Expression values were obtained from 5–9 biological replicates (pools of 2 to 4 youngest unfolded leaves at time of ASM or water treatment) from a single trial. **(b)** Relative expression of *MdAGG*, *MdPR2* and *MdPR10* from two independent experiments performed in the same conditions as in Fig. 5a (3-d heatwave, GDDH13 grafted plants – see afterwards). The youngest unfolded leaf at time of sampling was collected from 5 plants per experiment. **(c)** Protein immunodetection of defense-associated proteins. Equal quantities (10 µg) of each protein sample were loaded and run in three separate SDS-PAGE gels. Following transfer to PVDF membranes, each blot was probed with the appropriate primary antibody to detect MdAGG, MdPR2, and MdPR10. Each sample corresponds to a pool of proteins extracted from the 5 biological replicates per experiment presented in **(b)**.

As translation can be poorly correlated with transcription during immune regulation (Xu et al., 2017), we reasoned that high temperature may impede the translation of ASM-induced transcripts. However, western blots performed on *MdAGG*, *MdPR2* and *MdPR10*, three defense markers that had been previously associated with induced resistance at control temperature (Chavonet *et al*., 2022), revealed that protein accumulation was consistent with gene expression (Fig. 4b and c). Overall, the loss of ASM efficiency at high temperature was not captured by the variation of well-established defense markers, either at the transcript or protein level.

### High temperature dampens transcriptomic response to ASM but does not repress global defense

To explore the broader changes in the transcriptional landscape induced by ASM and high temperature, we performed a microarray analysis at the time of inoculation following a three-day heatwave (Fig. 5a). For this purpose, we took advantage of the ‘Golden Delicious’ doubled-haploid GDDH13 – the reference, homozygous apple genome (Daccord *et al*., 2017). We checked in GDDH13 grafted plants that high temperature also induced a loss of ASM efficiency (Fig. S4), which was broadly similar to, yet less severe than, the one observed in the seedlings of the ‘Golden Delicious’ progeny (Fig. 2b) – thereby suggesting that genetics is not the major source of variability in our pathosystem. We identified 3108 genes differentially expressed in at least one comparison (ASM vs. water at a given temperature, or 35°C vs. 21°C for a given ASM or water treatment). Overall, there was no strong transcriptomic effect of the prior heatwave for water-treated plants (Fig. 5b). Comparatively, ASM triggered massive transcriptomic changes, with patterns of up- and down-regulation that were largely conserved at both temperatures, yet the magnitude of these changes compared to their respective water controls was globally dampened at high temperature (Fig. 5b). GSEA analysis showed that ASM triggered global up-regulation in processes such as response to biotic stimulus, response to stress, signal transduction or alcohol metabolism, and down-regulation in processes such as cell division, photosynthesis, lipid metabolism, cell wall organization or response to hormones – at both temperatures (Fig. 5c). Interestingly, the functional categories for which up- or down-regulation was most dampened at high temperature were not focused on defense, but on cell division, signal transduction and various metabolic processes (Fig. 5c). Overall, ASM induced large transcriptomic regulation that was softened at high temperatures for some biological processes not obviously associated with defense.

**Fig. 5.**
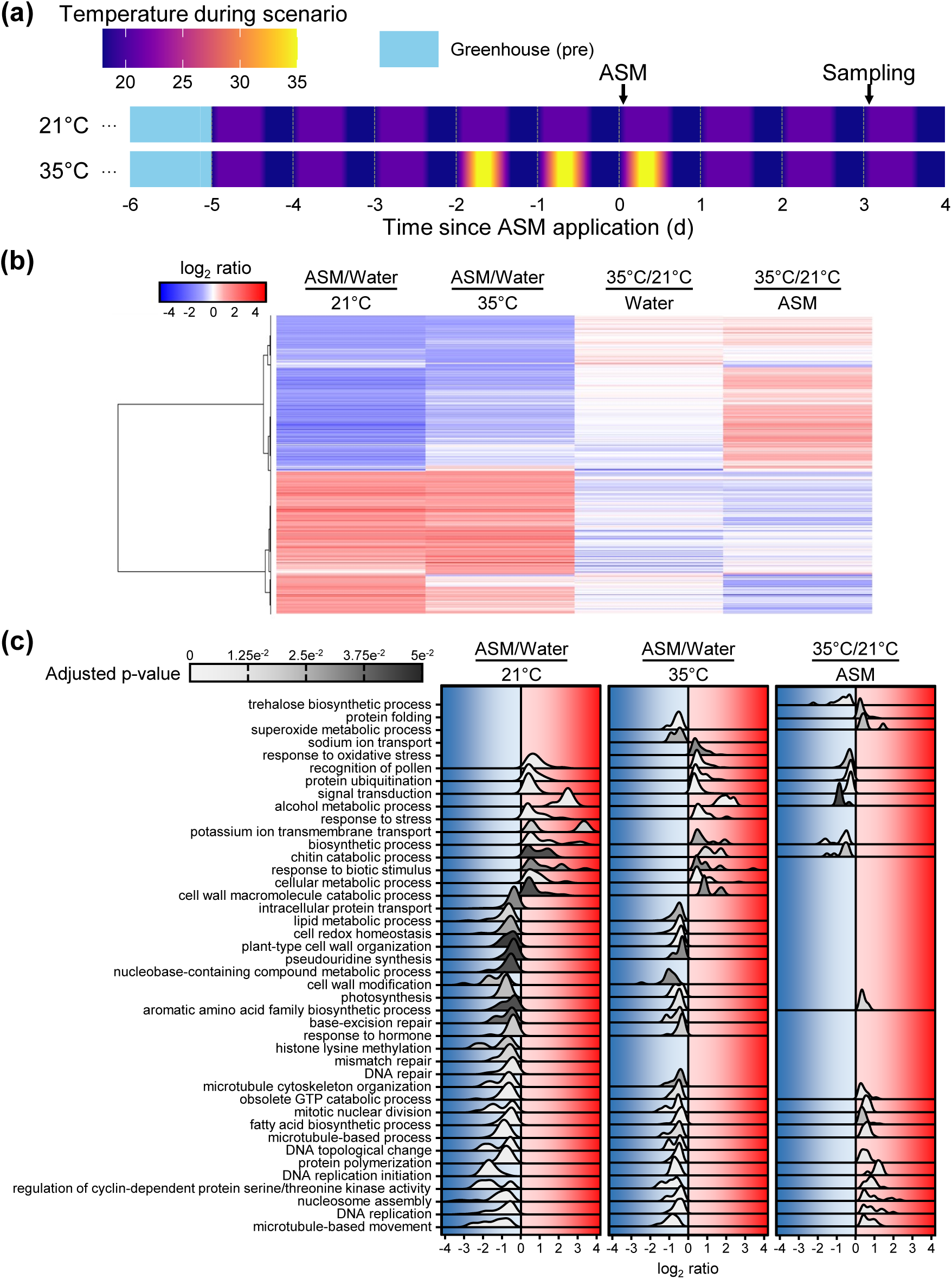
High temperature dampens transcriptomic response to ASM but does not repress global defense. **(a)** Schematic diagram of the experimental protocol. Air temperature during the contrasting thermal scenario (period in the growth chamber) is color-coded according to the color map. Arrows indicate the application of ASM or water treatment and the sampling of the youngest unfolded leaf of GDDH13 grafted plants dedicated to microarray analysis. **(b)** Heatmap of 3108 DEGs identified in at least one microarray comparison. DEGs were defined by a |log_2_ ratio| ≥ 1 and *P* < 0.05. The hierarchical clustering dendrogram was constructed from the dissimilarity matrix. Expression values were obtained from 2 biological replicates (each pooled from the youngest fully expanded leaf of 4 grafted plants) from 2 independent trials. **(c)** Ridgeline plot of GSEA on biological process ontology performed on comparison ASM/water for plants exposed to both 21°C and 35°C and 35°C/21°C comparison for ASM-treated plants. GSEA was obtained by 10.000 permutations with Benjamini and Hochberg procedure for p-value adjustment.

### Dampening of ASM-induced transcriptional regulation at high temperature is partial but pervasive

To confirm the transcriptional signature of the ASM × temperature interaction, we selected a set of representative genes for further RT-qPCR analyses on independent experiments. For this purpose, we first performed Clustering Affinity Search Technique (CAST) on the expression profiles of the 3108 genes that were differentially expressed in at least one comparison, generating 26 clusters (Fig. S5; Table S1). Among those, five clusters (Fig. 6) were highlighted as they were reminiscent of reduced resistance (R) or increased susceptibility (S) upon ASM application at high temperature, consistent with the phenotypes. Two clusters gathered the majority of genes and reflected the previously described global pattern, with an up- (cluster R1, 1130 genes) or down- (cluster S1, 1223 genes) regulation by ASM that is more or less dampened by high temperature. Such heat-induced dampening was exacerbated in clusters R2 (187 genes) and S2 (102 genes), respectively. Moreover, cluster R3 grouped 20 genes that were up-regulated by ASM at control temperature but down-regulated by ASM at high temperature (R3). Overall, clusters R1, R2 and R3 may be viewed as reservoirs of “resistance-associated genes” expected to be up-regulated by ASM, while clusters S1 and S2 may contain “susceptibility-associated genes” expected to be down-regulated by ASM – such up- or down-regulation being potentially vulnerable to high temperature, especially in clusters R2, R3 and S2. Supporting this classification, several PR-protein encoding genes were identified within cluster R1, whereas two genes previously implicated in apple susceptibility to *E. amylovora* (*DIPM3a* and *DIPM3b*; Meng *et al*., 2006) were identified within cluster S1 (Table S1).

**Fig. 6.**
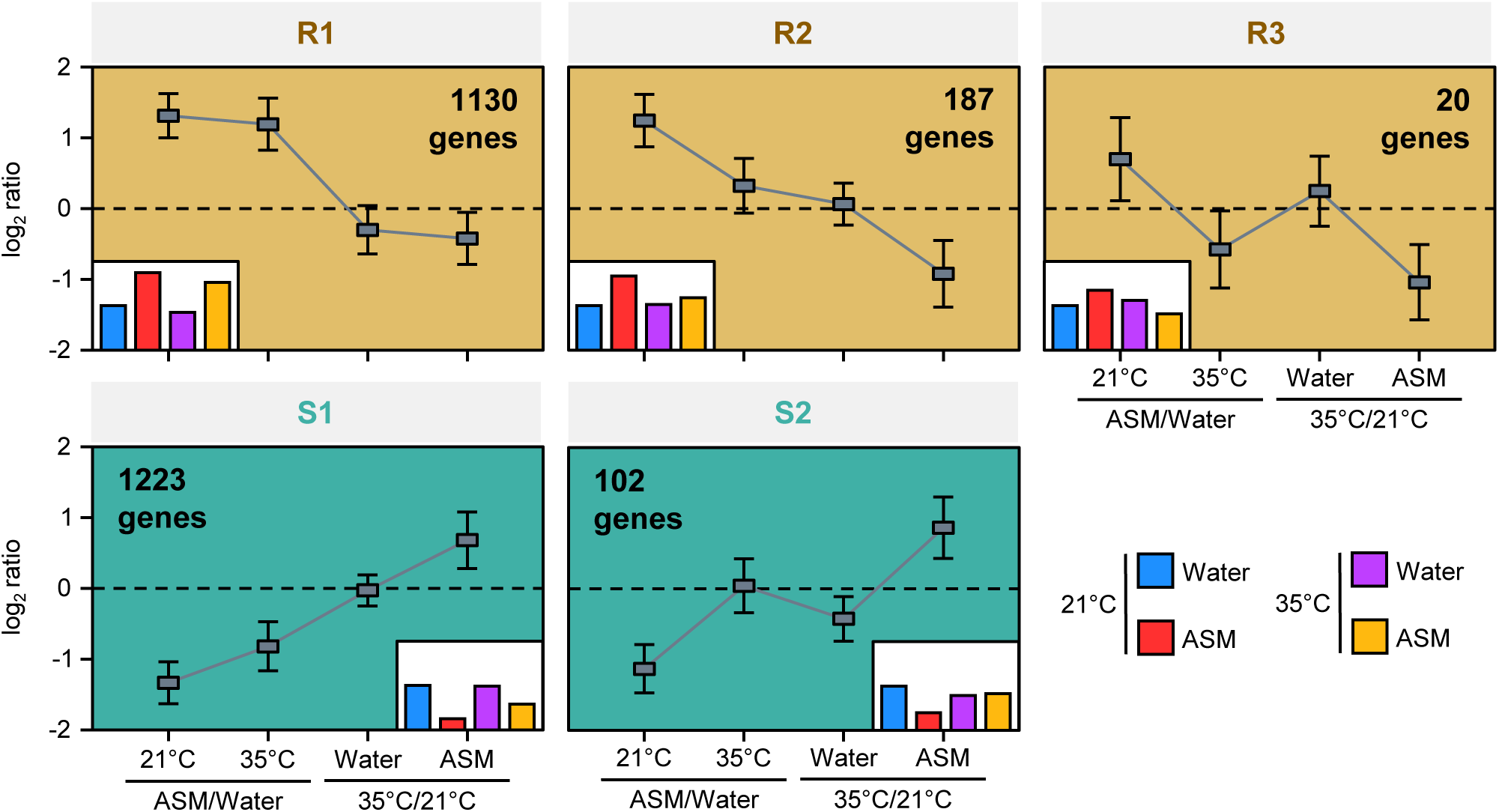
Identification of “resistance-associated” and “susceptibility-associated” marker genes by clustering. CAST clustering performed on the 3108 DEGs (Table S1 and Fig. S5) yielded several clusters evocative of resistant (R1, R2 and R3) and susceptible (S1 and S2) marker genes. The average pattern of expression ratio (± sd) is shown for each cluster. The number of DEGs in each resistance or susceptibility cluster are indicated in the upper right- or left-hand corner, respectively. To facilitate the interpretation of the variations in ratios, barplots showing the inferred expression pattern relative to the control (water, 21°C) are placed in the lower left- or right-hand corner of resistance and susceptibility clusters, respectively.

We selected 44 genes based on their distribution within these clusters and their functional annotation, including genes putatively involved in defense processes, but also in metabolism, cell cycle, transcription regulation, hormonal synthesis or cell wall properties (Table S2). We designed primers that targeted these genes and also potentially their homologues that were occasionally found across multiple clusters. As controls, we added the three defense genes known for their strong up-regulation by ASM and monitored previously: *MdAGG*, *MdPR2* and *MdPR10*. We then performed qPCR analyses that globally confirmed the expected expression patterns, with a large effect of ASM and a smaller effect of temperature (Fig. 7a). Consistent with this view, a PCA clearly separated the genes based on their belonging to different cluster types (“resistance-associated” *vs*. “susceptibility-associated”), and highlighted the major effect of ASM on gene expression (Fig. 7b). While only a few genes showed significant ASM × temperature interaction (Fig. 7a), whereby high temperature restricts the up- or down-regulation induced by ASM, the global interaction was evident on the PCA (Fig. 7b). This analysis confirms the partial but broad-spectrum erosion, at high temperature, of ASM efficiency for transcription regulation.

**Fig. 7.**
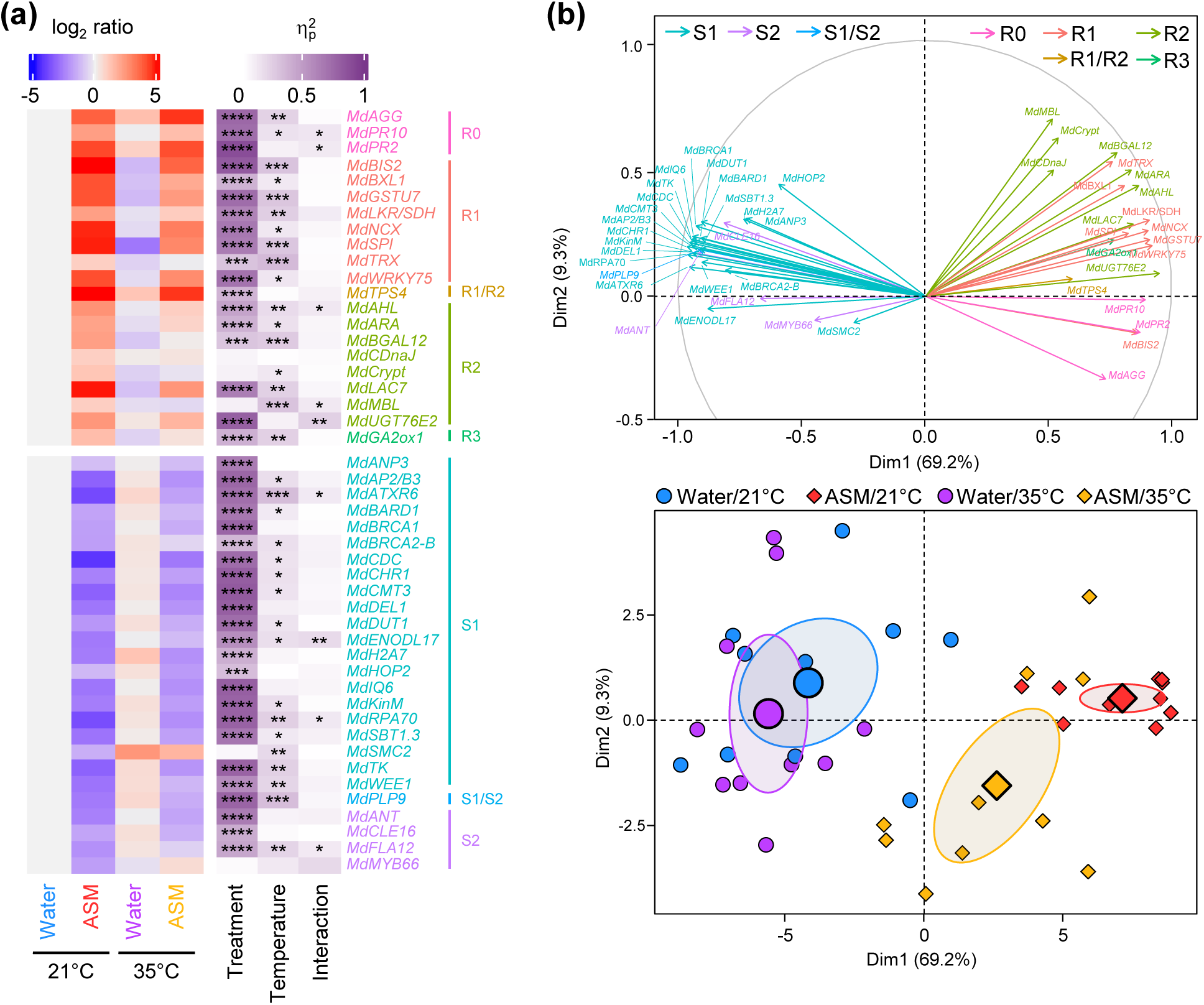
High temperature erodes ASM-induced transcriptional regulation of resistance- and susceptibility-associated marker genes. Expression values of water-treated leaves from plants exposed to 21°C were used for the calculation of the log_2_ ratio. Expression values were obtained from 10 biological replicates equally from 2 independent trials. **(a)** Heatmap showing relative gene expression and ANOVA effect sizes (η^2^_p_) of treatment, temperature and their interaction, as well as their respective significance level (*P<0.05, **P<0.01, ***P<0.001, ****P<0.0001). **(b)** PCA representation of relative gene expression on the two first dimensions, with the contribution of markers color-coded according to their cluster (top) and the distribution of samples grouped by treatment and temperature (bottom). Ellipses correspond to 95% confidence intervals around the centroid. Note that a single marker may be assigned to several clusters if the corresponding primer targets several homologous genes (Table S2).

### Transcription regulation by ASM is more vulnerable to posterior heatwaves

To further assess the vulnerability of ASM-induced transcriptional regulation to high temperature, we varied the number of days (1 to 4) of plant exposure to heatwave, and selected a subset of contrasting genes for qPCR analysis. The patterns of heat application were designed so that i) ASM or water treatments were always applied at the beginning of the last day of heatwaves, and ii) leaves were harvested at the same calendar time, 3 d after the ASM/water treatment (Fig. 8a). The 3-d heatwave corresponded to the same thermal regime as in the previous experiment, and the associated water treatment was therefore selected for data calibration. Overall, the effect of ASM was larger than that of the heatwave duration, and their interaction was minor (Fig. 8b). There was no obvious dose-effect of the duration, which was difficult to interpret. Nonetheless, the transcriptional effect of ASM towards “resistance-associated” markers (or away from “susceptibility-associated” markers) was maximal for the 1-d heatwave, while the amplitude of response to ASM was minimal for the 4-d heatwave (Fig. 8c). Thus, prolonged exposure to heatwave tended to increase the vulnerability of ASM-induced transcriptional regulation, yet the markers were not exquisitely sensitive to the heatwave duration when the heatwave was applied before ASM application.

**Fig. 8.**
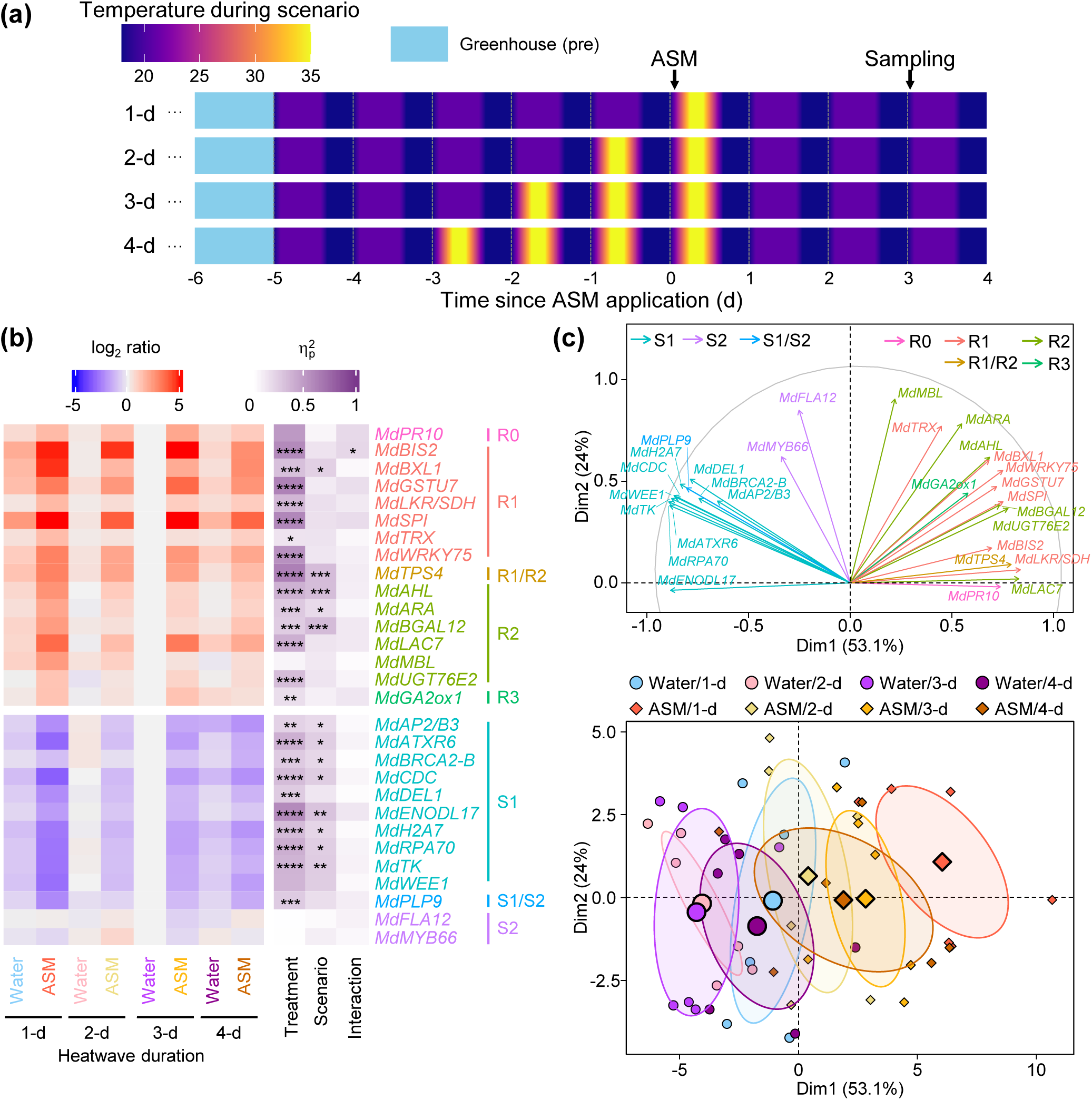
The duration of heat exposure before ASM treatment moderately affects the dampening of ASM-induced transcriptional regulation. Effect of heat duration prior to treatment on the expression of selected markers 3 d after ASM or water treatment, in grafted plants subjected to a 1-d, 2-d, 3-d and 4-d heatwave at 35°C. Expression values of water-treated leaves from plants exposed to 3-d 35°C were used for the calculation of the log_2_ ratio. Expression values were obtained from 6 biological replicates equally from 2 independent trials. **(a)** Schematic diagram of the experimental protocol. Air temperature during the contrasting thermal scenario (period in the growth chamber) is color-coded according to the color map. Arrows indicate the application of ASM (or water) treatment and the sampling of the youngest unfolded leaves. **(b)** Heatmap showing relative gene expression and ANOVA effect sizes (η^2^_p_) of treatment, temperature and their interaction, as well as their respective significance level (*P<0.05, **P<0.01, ***P<0.001, ****P<0.0001). **(c)** PCA representation of relative gene expression on the two first dimensions, with the contribution of markers color-coded according to their cluster (top) and the distribution of samples grouped by treatment and temperature (bottom). Ellipses correspond to 95% confidence intervals around the centroid. Note that a single marker may be assigned to several clusters if the corresponding primer targets several homologous genes (Table S2).

Finally, we varied the positioning of the heatwave in relation to the ASM application. A 3-d heatwave was imposed just before, during (as in the previous experiments) or just after the treatment application. Again, all GDDH13 leaves were harvested at the same calendar time, 3 d after the ASM/water treatment (Fig. 9a). Gene up- or down-regulation by ASM was more effective when the heatwave occurred before ASM application, and less effective when it occurred afterward, as confirmed by significant interaction between ASM and positioning (Fig. 9b) that was also evident in the PCA (Fig. 9c). A protection experiment on seedlings against *E. amylovora* corroborated that ASM efficiency is higher for a prior heatwave than for a posterior heatwave: when fire blight symptoms were scored at 7, 14 or 21 dpi, the p-value of the ASM effect (Fisher’s exact test within each condition) was always lower and around the significance threshold for a heatwave applied before ASM (Fig. S6). These results suggest that high temperature does not prime apple plants towards the erosion of ASM efficiency, but prevents ASM from mounting its full transcriptional regulation straight away.

**Fig. 9.**
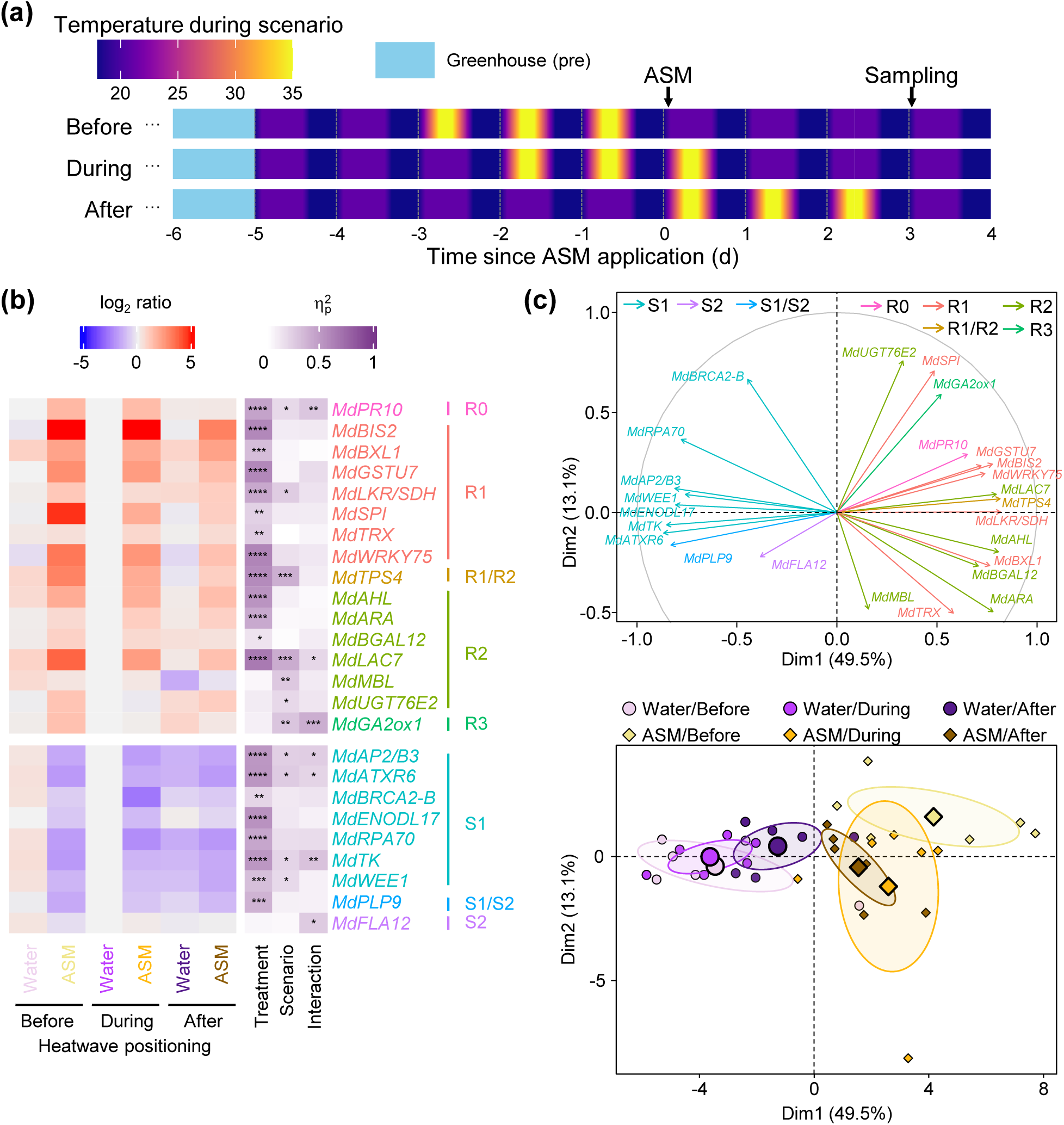
A heatwave posterior to ASM application dampens ASM-induced transcriptional regulation more strongly than a prior heatwave. Effect of heatwave positioning on expression of selected markers 3 d after ASM or water treatment, in grafted plants subjected to a 3-d heatwave at 35°C before, during or after ASM or water treatment. Expression values of water-treated leaves from plants under heatwave during the treatment were used for the calculation of the log_2_ ratio. Expression values were obtained from 6 biological replicates equally from 2 independent trials. **(a)** Schematic diagram of the experimental protocol. Air temperature during the contrasting thermal scenario (period in the growth chamber) is color-coded according to the color map. Arrows indicate the application of ASM (or water) treatment and the sampling of the youngest unfolded leaves. **(b)** Heatmap showing relative gene expression and ANOVA effect sizes (η^2^_p_) of treatment, temperature and their interaction, as well as their respective significance level (*P<0.05, **P<0.01, ***P<0.001, ****P<0.0001). **(c)** PCA representation of relative gene expression on the two first dimensions, with the contribution of markers color-coded according to their cluster (top) and distribution of samples grouped by treatment and temperature (bottom). Ellipses correspond to 95% confidence intervals around the centroid. Note that a single marker may be assigned to several clusters if the corresponding primer targets several homologous genes (Table S2).

## Discussion

Heatwaves are expected to become more prevalent with climate change and this may affect plant immunity and pathogen virulence, with positive or negative impacts on disease outcome depending on the pathosystem and the thermal scenario (Velásquez *et al*., 2018; Desaint *et al*., 2021; Chaloner *et al*., 2021; Roussin-Léveillée *et al*., 2024). Here, we found that subjecting apple plants to a short heatwave prior to infection with the bacteria *E. amylovora* or the fungus *V. inaequalis* does not impact the development of disease symptoms, but prevents ASM to drive an effective protection despite early accumulation of acibenzolar and active SA metabolism. Our results suggest that pre-exposure to high temperature affects ASM-induced immunity in apple plants, by preventing ASM from fully regulating a wide array of genes that are not solely related to defense.

Vulnerability of immunity at high temperature in Arabidopsis is underpinned by a temperature-susceptible module that normally drives the accumulation of SA and PR1 defense protein; however, ASM properly initiates NPR1-mediated signaling, and activates other gene modules to drive effective resistance upon warm conditions (Huot *et al*., 2017; Kim *et al*., 2022). In apple plants, we did not identify a major SA module vulnerable to high temperature, but rather found that ASM-induced gene regulation was globally less efficient following exposure to high temperature. Arabidopsis and other Brassicaceae mainly synthesize SA through the isochorismate synthase (ICS) pathway that has evolved in this family (Hong *et al*., 2025), while most other plants would rely on the phenylalanine ammonia-lyase (PAL) pathway (Hyun & Yoo, 2025). It is therefore plausible that thermovulnerability of SA metabolism is specific to Bassicaceae’s ICS pathway. The PAL pathway has been very recently elucidated (Liu *et al*., 2025; Wang *et al*., 2025; Zhu *et al*., 2025). Using our microarray data, we checked the pattern of the putative apple orthologues of the newly identified genes, and found that ASM upregulates the expression of several ones, especially at high temperature (Fig. S7), potentially explaining enhanced salicylates in these conditions (Fig. 3c-e) – but obviously not the loss of ASM-induced protection. Beyond SA metabolism, ASM normally rewires gene transcription so that defense responses are up-regulated while growth-associated processes are down-regulated (Huot *et al*., 2017; Warneys *et al*., 2018), which we confirmed here. At high temperature, ASM triggered changes in the transcriptome that were essentially similar to the ones at control temperature, yet attenuated. Overall, plants exposed to high temperature and treated with ASM showed a transcriptomic signature evocative of lower defense and higher susceptibility compared to ASM-treated plants at control temperature (Fig. 10). The mechanisms by which high temperature eventually suppresses ASM-induced protection in apple plants may therefore involve many end-players that each partly contributes to plant resistance or susceptibility. For instance, heat stress partially impaired ASM-induction of downstream resistance end-players with established roles against different pests. These included e.g. *PR-10* (*PATHOGENESIS RELATED-10*), homologues of ribonuclease proteins that possess antimicrobial properties (Liu & Ekramoddoullah, 2006); *BIS2* (*BIPHENYL SYNTHASE 2*), enzymes involved in biosynthesis of biphenyl and dibenzofuran phytoalexins in apple shoots, with antimicrobial activity against *E. amylovora* (Chizzali *et al*., 2012) and *V. inaequalis* (Hrazdina *et al*., 1997); or *TPS4* (*TERPENE SYNTHASE 4*), enzymes that participate in the synthesis of defensive volatile compounds in apple (Nieuwenhuizen *et al*., 2013) such as (*E*,*E*)-*α*-farnesene, a sesquiterpene repellent against rosy apple aphid (*Dysaphis plantaginea*; Warneys *et al*., 2018). Thus, ASM-induced resistance was transcriptionally compromised by heat, correlating here with a loss of protection against *E. amylovora* and *V. inaequalis*.

**Fig. 10.**
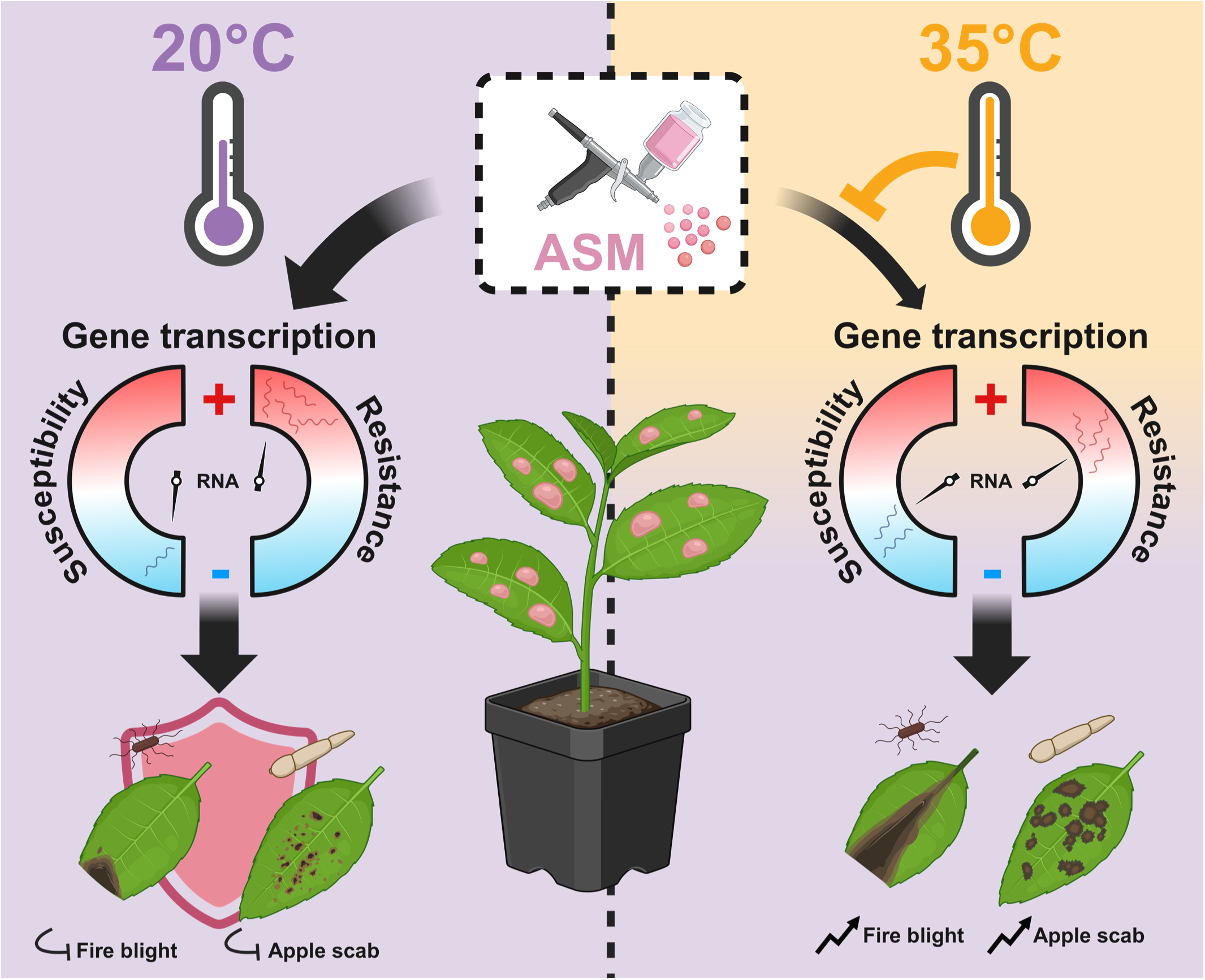
Working model for the loss of ASM-induced protection of apple plants upon transient heat. At control temperature (20°C), spraying apple plants with ASM prior to inoculation with the bacteria *E. amylovora* or the fungus *V. inaequalis* allows apple leaves to mount an effective transcriptional response whereby resistance-associated genes are up-regulated and susceptibility-associated genes are down-regulated – which reduces the incidence of fire blight or apple scab. When a heatwave (35°C) occurs in between ASM application and pathogen inoculation, ASM-induced transcriptional regulation is dampened, ultimately enabling pathogens to proliferate. Created in BioRender.com.

Conversely, ASM may normally repress susceptibility or *S* genes that facilitate compatibility with the pathogen, and fails to do so after heat stress, resulting in enhanced plant susceptibility. Only a handful of susceptibility genes are known for the apple / *E. amylovora* pathosystem, encoding apple proteins that interact with primary pathogenicity effectors (Campa *et al*., 2019; Pompili *et al*., 2020; Tegtmeier *et al*., 2020), while none have been formally described for *V. inaequalis* so far. Our transcriptomic results support the view that heat stress dampens ASM-driven down-regulation of some of these *S* genes. Further *S* genes likely remain to be discovered, including those related to host physiology that may be required to allow sustained compatibility, as exemplified in other pathosystems (van Schie & Takken, 2014). While our “susceptibility-associated” markers cannot be defined as genuine *S* genes, their de-repression upon heat may enhance plant susceptibility. These markers contained a number of genes plausibly associated with DNA replication or integrity – based on their Arabidopsis homologs (i.e. *WEE1*, *RPA70*, *TK*, *BRCA2*, *ATXR6* – see Table S2 for full gene names). Other markers were possibly linked to cell wall properties (*FLA12*; MacMillan *et al*., 2010), solute transport (*ENODL17*; Denancé *et al*., 2014), or cell death execution (*PLP9*; La Camera *et al*., 2005, 2009), providing a diversity of potential entries whereby pathogens may divert host physiology. In this framework, the erosion of ASM-induced down-regulation of genes useful to plant physiology under heat, combined with the reduced induction of defense mechanisms, would be exploited by the pathogen – effectively leveraging both sides of the “growth– defense tradeoff” (Huot *et al*., 2014; Monson *et al*., 2022).

High temperature seems therefore to directly impede the transcriptional responses to ASM. In line with this idea, the dampened transcriptional pattern was exacerbated when positioning a 3-d heatwave just after the ASM treatment, i.e. when ASM-induced transcription is fully active (Warneys *et al*., 2018; Chavonet *et al*., 2022). The molecular mechanisms that link high-temperature perception to transcriptional regulation are increasingly well-described, with the recent discovery of several plant thermosensors (Casal *et al*., 2024). Interestingly, the thermosensor THERMO-WITH ABA-RESPONSE 1 (TWA1) is a predicted intrinsically disordered protein that changes its conformation upon high temperature, allowing physical interaction with the JASMONATE-ASSOCIATED MYC2–LIKE 2 (JAM2) transcription factor and the TOPLESS (TPL) transcriptional repressor, which down-regulates the expression of target genes and optimizes heat stress response (Bohn *et al*., 2024). JAM2 is a negative regulator of JA signaling (Sasaki-Sekimoto *et al*., 2013), a hormonal pathway that is down-regulated by *E. amylovora* during the infection of susceptible apple plants (Dugé de Bernonville *et al*., 2012). Moreover, the classical JA–SA antagonism in Arabidopsis may turn into synergism under warm conditions (Roussin-Léveillée *et al*., 2024), due to a tunable regulation loop in the expression of SA biosynthesis genes by JA (Mine *et al*., 2017). This illustrates potential connections between high temperature sensing and response to biotic cues through transcriptional regulation. It should be stressed, however, that dampening of transcriptional regulation was far from complete, suggesting that the loss of ASM-induced protection at high temperature involves additional layers of biological modifications that are not fully captured by transcripts. Tracking ASM-induced changes in the metabolome upon high temperature could improve our understanding of the mechanisms underlying the loss of protection we report.

In conclusion, our study unveils a loss of ASM-induced protection against fire blight and apple scab upon transient exposure to heat stress. It covaries with a pervasive dampening of ASM-induced transcription – and probably further physiological rearrangements – so that high temperature makes ASM-treated plants globally less capable of producing defense-related compounds toxic to microbes, and more prone to accumulate susceptible factors that facilitate microbial colonization. As plant response to ASM involves the regulation of many biological processes beyond defense, the protection efficacy of ASM is shaped by the environment due to the interplay between defense and physiology in plants. Disentangling this relationship would enhance our basic understanding of plant immunity in a changing climate, and may offer prospects for stabilizing the efficacy of plant resistance inducers in the field.

## Acknowledgments

We thank Anthony Juillard, Brice Marolleau and Mylène Ruh for their involvement in disease assessment assays, the platforms ANAN (analysis of nucleic acids), PHYTO (Phytochemical analyses) and PHENOTIC (Angers Plant Phenotyping Facility, https://doi.org/10.17180/YKBZ-2V85) of the SFR QUASAV for excellent technical assistance, and the UE Horti (https://doi.org/10.15454/1.5573931618268674E12) for the supply of scions. This research was conducted in the framework of i) the regional program “Objectif Végétal, Research, Education and Innovation in Pays de la Loire”, supported by the French Region Pays de la Loire, Angers Loire Métropole and the European Regional Development Fund, ii) the France Agrimer Connaissances Program managed by the French Ministry of Agriculture (Project FAM-Connaissances 2023-11429220-OCASE) and iii) the French Priority Research Programme “Cultiver et Protéger Autrement” (PPR-CPA) managed by the Agence Nationale de la Recherche (project ANR-20-PCPA-0003 CapZeroPhyto). This work was also supported by the Czech Science Foundation (GACR) project No. 22-17435S, and by grants from the Ministère de l’Education Nationale, de l’Enseignement Supérieur et de la Recherche, France (to E.C.) and from INRAE-AGROECOSYSTEM Department and the Region Pays de la Loire, France (to B-h.N.). B-h.N. received a travel grant from VAAME Doctoral School for a stay in the Laboratory of Growth Regulators.

## Author contributions

A.D., F.P., M.G., R.L. and M.-N.B. conceived the project and designed the research plan. E.C. and B.-h.N. performed most of the experiments, with the contribution of R.C. and C.H. for some experiments. J.Š. and O.N. supervised hormonal analyses. A.D., F.P., M.G., R.L., M.-N.B., E.C. and B.-h.N. analyzed the data and wrote the manuscript. All authors read and approved the final manuscript.

## Data availability

The microarray data sets presented in this study can be found in online repositories. The names of the repository and accession number can be found at: https://www.ncbi.nlm.nih.gov/geo/query/acc.cgi?acc=GSE280234.

## Supporting Information

Additional Supporting Information may be found online in the Supporting Information section at the end of the article.

**Figure S1.** Climatic variables measured at the meteorological station of Beaucouzé (France) for the eleven hottest days recorded between March 2016 and July 2018.

**Figure S2.** Climate dynamics applied in the growth chamber to study the effect of heatwaves on apple response to ASM.

**Figure S3.** Impact of heat and treatment on plant phenotype, and representative photographs of apple seedlings and their typical disease symptoms.

**Figure S4.** Accumulation of high temperatures prior to inoculation impairs the ability of apple grafts of GDDH13 to mount an effective ASM-induced protection against *E. amylovora*.

**Figure S5.** Identification of gene clusters by CAST.

**Figure S6.** Protection provided by ASM to apple seedlings against *E. amylovora* is more strongly dampened by a heatwave posterior to ASM application than by a prior heatwave.

**Figure S7.** Putative apple genes of SA biosynthesis pathway are induced by ASM at high temperature.

**Table S1.** List of genes that are differentially regulated by ASM treatment and transient heat according to the microarray analysis.

**Table S2.** Design of “resistance” and “susceptibility” marker genes.

## Supplementary figures captions

**Fig. S1. Climatic variables measured at the meteorological station of Beaucouzé (France) for the eleven hottest days recorded between March 2016 and July 2018.** Empty circles correspond to individual, hourly values. **(a)** Air temperature, **(b)** relative humidity (RH), **(c)** solar radiation and corresponding percent irradiance. Grey solid curves represent the average trend as determined by the non-parametric loess method (span = 0.5), surrounded by the standard error (grey shaded area). Red dashed lines indicate the piecewise linear approximation of each climatic variable that was used to design the heatwave scenarios in the growth chamber.

**Fig. S2. Climate dynamics applied in the growth chamber to study the effect of heatwaves on apple response to ASM. (a)** “Control”, **(b)** “Short” and **(c)** “Long” scenarios showing the daily variations in temperature, relative humidity (RH) and percent irradiance applied in growth chambers. Combination of these elemental daily programs were used to reproduce the different thermal scenarios that were studied.

**Fig. S3. Impact of heat and treatment on plant phenotype, and representative photographs of apple seedlings and their typical disease symptoms. (a)**, **(b)**, and **(c)** Boxplots and jitter plots showing the impact of temperature scenario and ASM or water treatment on apple seedling phenotypes. All measurements were performed on the same plants of the same experiment (Methods). At time 0, plants were sprayed with ASM or water and immediately transferred from a greenhouse to a growth chamber with controlled environmental conditions. Air temperature was maintained at either 20°C or 35°C for 24 h, and all plants then remained in the growth chamber at 20°C during 48 h more. The elongation rate of **(a)** the stem and **(b)** the youngest unfolded leaf was calculated using measurements made before treatment and 3 d after treatment. Statistical differences between ASM and water treatment were determined at each temperature using pairwise t-tests adjusted by the Bonferroni method, and were deemed not significant (ns). **(c)** Stomatal conductance (g_s_) was measured on the youngest unfolded leaf at time of ASM/water treatment application (left panel) and on the leaf rank immediately below (right panel), 1.5 h before treatment and 3, 48 and 72 h after treatment. Multiple mean comparisons were performed between each combination of both factors at each time point using pairwise t-tests adjusted by the Bonferroni method (* 0.05≥P>0.01, ** 0.01≥*P*>0.001). Note that non-significant differences have been omitted for clarity. **(d)** Photograph of a representative batch of apple seedlings at the start of the experiment. **(e)** Photograph of systemic fire blight necrosis symptoms on a diseased apple seedling, 21 d after inoculation by *E. amylovora*. The inoculated leaf is indicated by a white arrowhead and the picture focuses on necrotic symptoms spreading along the midvein of a distant leaf, and along the stem up to the plant apex. **(f)** Photograph of an apple seedling leaf showing *V. inaequalis* sporulation, 21 d after inoculation.

**Fig. S4. Accumulation of high temperatures prior to inoculation impairs the ability of apple grafts of GDDH13 to mount an effective ASM-induced protection against *E. amylovora*. (a)** Schematic representation of the experimental protocol (details in Methods). Air temperature during the contrasting thermal scenario (period in the growth chamber) is color-coded according to the color map. Arrows indicate the application of ASM (or water) treatments and pathogen inoculation on the youngest unfolded leaf. **(b)** Stacked bar chart showing the proportion of healthy (asymptomatic on stem) and diseased (symptomatic on stem) grafted plants inoculated with *E. amylovora*, 28 d post-inoculation. Data represent 48 biological replicates from 4 independent trials (n = total number of plants). Statistical significance was determined using pairwise Fisher’s exact tests; same letters indicate groups that are not significantly different. Icons created in BioRender.com. **(c)** Photograph of a diseased GDDH13 graft presenting fire blight symptoms on the apex and displaying the typical “hook shape”.

**Fig. S5. Identification of gene clusters by CAST.** Clustering was based on the expression pattern of 3108 genes that displayed differential expression (defined by a |log_2_ ratio| ≥ 1 and P < 0.05 in at least one microarray comparison, see Fig. 5b) in plants treated with ASM or water and exposed to control (21°C) or elevated (35°C) temperature. Pearson correlation distance metric (0.95 threshold affinity) allowed the construction of 26 clusters. Grey lines correspond to the expression ratio pattern of each individual DEG. Red points correspond to the average log_2_ expression ratio (± sd) within each comparison, for each cluster. The number of DEGs grouped in each cluster and the corresponding cluster number (and name for R1, R2, R3, S1 and S2 investigated in Fig. 6) are indicated in the upper right- and left-hand corner of each box, respectively.

**Fig. S6. Protection provided by ASM to apple seedlings against *E. amylovora* is more strongly dampened by a heatwave posterior to ASM application than by a prior heatwave. (a)** Schematic representation of the experimental protocol (details in Methods). Air temperature during the contrasting thermal scenario (period in the growth chamber) is color-coded according to the color map. Arrows indicate the application of ASM (or water) treatments and pathogen inoculation on the youngest unfolded leaf. **(b)** Stacked bar charts showing the proportion of healthy (asymptomatic on stem) and diseased (symptomatic on stem) seedlings inoculated with *E. amylovora* at 7, 14 and 21 d post-inoculation (dpi). Data represent 55–56 biological replicates (number of plants in each condition) from a single trial. The *p*-value of Fisher’s exact test between water and ASM treatments is shown above each pair of barplot (^•^P<0.1, *P<0.05). Icons created in BioRender.com.

**Fig. S7. Putative apple genes of SA biosynthesis pathway are induced by ASM at high temperature.** The expression pattern of putative apple orthologues of genes involved in SA biosynthesis was extracted from microarray data on GDDH13 leaves (Fig. 5). For each enzymatic step, expression of corresponding gene(s) are color-coded in boxes showing the effect of ASM ① at 21°C and ② at 35°C, and the effect of high temperature upon treatment with ③ water and ④ ASM. Significant up- or down-regulations are notified (*). The gene represented by grey boxes was absent from the microarray. (a) Phenylalanine ammonia-lyase (PAL) pathway. Apple protein orthologues of cinnamoyl-coenzyme A ligase (CNL) / *Oryza sativa* SA-DEFICIENT GENE 1 (OSD1), ABNORMAL INFLORESCENCE MERISTEM1 (AIM1), 3-ketoacyl-CoA thiolase (KAT), benzyl alcohol benzoyltransferase (BEBT) / OSD2, benzyl benzoate oxidase (BBO) / ODS3 / benzylbenzoate hydroxylase (BBH) and benzyl salicylate hydrolase (BSH) / OSD4 / benzylsalicylate esterase (BSE) were identified according to a BLAST of protein sequence from rice. Apple protein with an e-value of 0, less than 10% of gaps and 60% of identities were retained. If no protein met these criteria, the top four hits were selected. (b) Isochorismate synthase (ICS) pathway. Apple mRNA orthologues of ICS, AVRPPHB SUSCEPTIBLE 3 (PBS3) and ENHANCED PSEUDOMONAS SUSCEPTIBILITY 1 (EPS1) were identified according to a BLAST with at least 60% of identities with CDS sequence from *Arabidopsis thaliana*.

## Supplementary tables captions

**Table S1. List of genes that are differentially regulated by ASM treatment and transient heat according to the microarray analysis.** Differentially expressed genes were defined by a |log_2_ ratio| ≥ 1 and P < 0.05. The table shows gene accession, gene description, gene symbol (if any), log_2_ fold change within each comparison, cluster membership (see Fig. S5), and functional annotation. Gene description and functional annotation are based on the GDDH13 genome annotation (Daccord *et al*., 2017).

**Table S2. Design of “resistance” and “susceptibility” marker genes.** Genes initially selected through CAST belonged to “resistance” (R1, R2, R3) or “susceptibility” (S1, S2) clusters (Fig. 6). Homologous genes were searched in the genome, and were eventually found in the same or an alternative cluster. Primers were designed to target as many homologous genes as possible, and the targeted ones are bolded. The table shows gene name, functional annotation, accession, cluster membership (if any), and primer sequence and concentration. Functional annotation is based on the GDDH13 genome annotation (Daccord *et al*., 2017).

## References

Aćimović SG, Zeng Q, McGhee GC, Sundin GW, Wise JC. 2015. Control of fire blight (*Erwinia amylovora*) on apple trees with trunk-injected plant resistance inducers and antibiotics and assessment of induction of pathogenesis-related protein genes. Frontiers in Plant Science 6: 16.

Ben-Dor A, Shamir R, Yakhini Z. 1999. Clustering gene expression patterns. Journal of Computational Biology 6: 281–297.

Bohn L, Huang J, Weidig S, Yang Z, Heidersberger C, Genty B, Falter-Braun P, Christmann A, Grill E. 2024. The temperature sensor TWA1 is required for thermotolerance in *Arabidopsis*. Nature 629: 1126– 1132.

Bowen JK, Mesarich CH, Bus VGM, Beresford RM, Plummer KM, Templeton MD. 2011. *Venturia inaequalis*: the causal agent of apple scab. Molecular Plant Pathology 12: 105–122.

Brisset M-N, Cesbron S, Thomson SV, Paulin J-P. 2000. Acibenzolar-S-methyl induces the accumulation of defense-related enzymes in apple and protects from fire blight. European Journal of Plant Pathology 106: 529–536.

Calenge F, Faure A, Goerre M, Gebhardt C, Van de Weg WE, Parisi L, Durel C-E. 2004. Quantitative trait loci (QTL) analysis reveals both broad-spectrum and isolate-specific QTL for scab resistance in an apple progeny challenged with eight isolates of *Venturia inaequalis*. Phytopathology 94: 370–379.

Campa M, Piazza S, Righetti L, Oh C-S, Conterno L, Borejsza-Wysocka E, Nagamangala KC, Beer SV, Aldwinckle HS, Malnoy M. 2019. *HIPM* is a susceptibility gene of *Malus* spp.: Reduced expression reduces susceptibility to *Erwinia amylovora*. Molecular Plant-Microbe Interactions 32: 167–175.

Canet JV, Dobón A, Ibáñez F, Perales L, Tornero P. 2010a. Resistance and biomass in Arabidopsis: a new model for Salicylic Acid perception. Plant Biotechnology Journal 8: 126–141.

Canet JV, Dobón A, Roig A, Tornero P. 2010b. Structure-function analysis of *npr1* alleles in Arabidopsis reveals a role for its paralogs in the perception of salicylic acid. *Plant*, Cell & Environment 33: 1911–1922.

Casal JJ, Murcia G, Bianchimano L. 2024. Plant thermosensors. Annual Review of Genetics 58: 135– 158.

Celton J-M, Gaillard S, Bruneau M, Pelletier S, Aubourg S, Martin-Magniette M-L, Navarro L, Laurens F, Renou J-P. 2014. Widespread anti-sense transcription in apple is correlated with siRNA production and indicates a large potential for transcriptional and/or post-transcriptional control. New Phytologist 203: 287– 299.

Chaloner TM, Gurr SJ, Bebber DP. 2021. Plant pathogen infection risk tracks global crop yields under climate change. Nature Climate Change 11: 710–715.

Chavonet E, Gaucher M, Warneys R, Bodelot A, Heintz C, Juillard A, Cournol R, Widmalm G, Bowen JK, Hamiaux C, et al. 2022. Search for host defense markers uncovers an apple agglutination factor corresponding with fire blight resistance. Plant Physiology 188: 1350–1368.

Cheng C, Gao X, Feng B, Sheen J, Shan L, He P. 2013. Plant immune response to pathogens differs with changing temperatures. Nature Communications 4: 2530.

Chizzali C, Gaid MM, Belkheir AK, Hänsch R, Richter K, Flachowsky H, Peil A, Hanke M-V, Liu B, Beerhues L. 2012. Differential expression of biphenyl synthase gene family members in fire-blight-infected apple ‘Holsteiner Cox’. Plant Physiology 158: 864–875.

Coupel-Ledru A, Westgeest AJ, Albasha R, Millan M, Pallas B, Doligez A, Flutre T, Segura V, This P, Torregrosa L, et al. 2024. Clusters of grapevine genes for a burning world. New Phytologist 242: 10–18.

Daccord N, Celton J-M, Linsmith G, Becker C, Choisne N, Schijlen E, van de Geest H, Bianco L, Micheletti D, Velasco R, et al. 2017. High-quality *de novo* assembly of the apple genome and methylome dynamics of early fruit development. Nature Genetics 49: 1099–1106.

Denancé N, Szurek B, Noël LD. 2014. Emerging functions of nodulin-like proteins in non-nodulating plant species. Plant and Cell Physiology 55: 469–474.

Desaint H, Aoun N, Deslandes L, Vailleau F, Roux F, Berthomé R. 2021. Fight hard or die trying: when plants face pathogens under heat stress. New Phytologist 229: 712–734.

Ding Y, Sun T, Ao K, Peng Y, Zhang Y, Li X, Zhang Y. 2018. Opposite roles of salicylic acid receptors NPR1 and NPR3/NPR4 in transcriptional regulation of plant immunity. Cell 173: 1454–1467.e15.

Dugé de Bernonville T, Gaucher M, Flors V, Gaillard S, Paulin J-P, Dat JF, Brisset M-N. 2012. T3SS-dependent differential modulations of the jasmonic acid pathway in susceptible and resistant genotypes of *Malus* spp. challenged with *Erwinia amylovora*. Plant Science 188–189: 1–9.

Dugé de Bernonville T, Marolleau B, Staub J, Gaucher M, Brisset M-N. 2014. Using molecular tools to decipher the complex world of plant resistance inducers: an apple case study. Journal of Agricultural and Food Chemistry 62: 11403–11411.

Eckardt NA. 2002. Plant disease susceptibility genes? The Plant Cell 14: 1983–1986.

Floková K, Tarkowská D, Miersch O, Strnad M, Wasternack C, Novák O. 2014. UHPLC–MS/MS based target profiling of stress-induced phytohormones. Phytochemistry 105: 147–157.

Friedrich L, Lawton K, Ruess W, Masner P, Specker N, Gut Rella M, Meier B, Dincher S, Staub T, Uknes S, et al. 1996. A benzothiadiazole derivative induces systemic acquired resistance in tobacco. The Plant Journal 10: 61–70.

Gessler C, Patocchi A, Sansavini S, Tartarini S, Gianfranceschi L. 2006. *Venturia inaequalis* resistance in apple. Critical Reviews in Plant Sciences 25: 473–503.

Görlach J, Volrath S, Knauf-Beiter G, Hengy G, Beckhove U, Kogel K-H, Oostendorp M, Staub T, Ward E, Kessmann H, et al. 1996. Benzothiadiazole, a novel class of inducers of systemic acquired resistance, activates gene expression and disease resistance in wheat. The Plant Cell 8: 629–643.

Gorshkov V, Tsers I. 2022. Plant susceptible responses: the underestimated side of plant–pathogen interactions. Biological Reviews 97: 45–66.

Gu Z. 2022.Complex heatmap visualization. iMeta 1: e43.

Guillaumès J, Chevalier M, Parisi L. 1995. Étude des relations *Venturia inaequalis* — *Malus* × *domestica* sur vitroplants. Revue Canadienne de Phytopathologie 17: 305–311.

Holb IJ. 2007. Classification of apple cultivar reactions to scab in integrated and organic production systems. Canadian Journal of Plant Pathology 29: 251–260.

Hong K, Nakano M, Tang Y, Jeanguenin L, Kang W, Wang Y, Zuo L, Li P, He J, Jiang W, et al. 2025. Emergence of isochorismate-based salicylic acid biosynthesis within Brassicales. Proceedings of the National Academy of Sciences of the United States of America 122: e2506170122.

Hrazdina G, Borejsza-Wysocki W, Lester C. 1997. Phytoalexin production in an apple cultivar resistant to *Venturia inaequalis*. Phytopathology 87: 868–876.

Hung J-H, Yang T-H, Hu Z, Weng Z, DeLisi C. 2012. Gene set enrichment analysis: performance evaluation and usage guidelines. Briefings in Bioinformatics 13: 281–291.

Huot B, Castroverde CDM, Velásquez AC, Hubbard E, Pulman JA, Yao J, Childs KL, Tsuda K, Montgomery BL, He SY. 2017. Dual impact of elevated temperature on plant defence and bacterial virulence in *Arabidopsis*. Nature Communications 8: 1808.

Huot B, Yao J, Montgomery BL, He SY. 2014. Growth–defense tradeoffs in plants: a balancing act to optimize fitness. Molecular Plant 7: 1267–1287.

Hyun J, Yoo H. 2025. Missing steps uncovered in a pathway plants use to produce the defence molecule salicylic acid. Nature 645: 48–50.

Kassambara A. 2023a. rstatix: Pipe-friendly framework for basic statistical tests. R package version 0.7.2.

Kassambara A. 2023b. ggpubr: ‘ggplot2’ based publication ready plots. R package version 0.6.0.

Kassambara A, Mundt F. 2020. factoextra: Extract and visualize the results of multivariate data analyses. R package version 1.0.7.

Kim JH, Castroverde CDM, Huang S, Li C, Hilleary R, Seroka A, Sohrabi R, Medina-Yerena D, Huot B, Wang J, et al. 2022. Increasing the resilience of plant immunity to a warming climate. Nature 607: 339– 344.

King EO, Ward MK, Raney DE. 1954. Two simple media for the demonstration of pyocyanin and fluorescin. Journal of Laboratory and Clinical Medicine 44: 301–307.

Kostick SA, Norelli JL, Evans KM. 2019. Novel metrics to classify fire blight resistance of 94 apple cultivars. Plant Pathology 68: 985–996.

Kuźniak E, Głowacki R, Chwatko G, Kopczewski T, Wielanek M, Gajewska E, Skłodowska M. 2014. Involvement of ascorbate, glutathione, protein S-thiolation and salicylic acid in benzothiadiazole-inducible defence response of cucumber against *Pseudomonas syringae* pv *lachrymans*. Physiological and Molecular Plant Pathology 86: 89–97.

La Camera S, Balagué C, Göbel C, Geoffroy P, Legrand M, Feussner I, Roby D, Heitz T. 2009. The *Arabidopsis* patatin-like protein 2 (PLP2) plays an essential role in cell death execution and differentially affects biosynthesis of oxylipins and resistance to pathogens. Molecular Plant-Microbe Interactions 22: 469– 481.

La Camera S, Geoffroy P, Samaha H, Ndiaye A, Rahim G, Legrand M, Heitz T. 2005. A pathogen-inducible patatin-like lipid acyl hydrolase facilitates fungal and bacterial host colonization in Arabidopsis. The Plant Journal 44: 810–825.

Lawton KA, Friedrich L, Hunt M, Weymann K, Delaney T, Kessmann H, Staub T, Ryals J. 1996. Benzothiadiazole induces disease resistance in *Arabidopsis* by activation of the systemic acquired resistance signal transduction pathway. The Plant Journal 10: 71–82.

Lê S, Josse J, Husson F. 2008. FactoMineR: An R package for multivariate analysis. Journal of Statistical Software 25: 1–18.

Liu J-J, Ekramoddoullah AKM. 2006. The family 10 of plant pathogenesis-related proteins: Their structure, regulation, and function in response to biotic and abiotic stresses. Physiological and Molecular Plant Pathology 68: 3–13.

Liu Y, Xu L, Wu M, Wang J, Qiu D, Lan J, Lu J, Zhang Y, Li X, Zhang Y. 2025. Three-step biosynthesis of salicylic acid from benzoyl-CoA in plants. Nature 645: 201–207.

Long SP, Ort DR. 2010. More than taking the heat: crops and global change. Current Opinion in Plant Biology 13: 240–247.

MacMillan CP, Mansfield SD, Stachurski ZH, Evans R, Southerton SG. 2010. Fasciclin-like arabinogalactan proteins: specialization for stem biomechanics and cell wall architecture in Arabidopsis and *Eucalyptus*. The Plant Journal 62: 689–703.

Malamy J, Hennig J, Klessig DF. 1992. Temperature-dependent induction of salicylic acid and its conjugates during the resistance response to tobacco mosaic virus infection. The Plant Cell 4: 359–366.

Mangiafico SS. 2024. rcompanion: Functions to support extension education program evaluation. R package version 2.4.36.

Marchin RM, Backes D, Ossola A, Leishman MR, Tjoelker MG, Ellsworth DS. 2022. Extreme heat increases stomatal conductance and drought-induced mortality risk in vulnerable plant species. Global Change Biology 28: 1133–1146.

Marolleau B, Gaucher M, Heintz C, Degrave A, Warneys R, Orain G, Lemarquand A, Brisset M-N. 2017. When a plant resistance inducer leaves the lab for the field: integrating ASM into routine apple protection practices. Frontiers in Plant Science 8: 1938.

Maxson-Stein K, He S-Y, Hammerschmidt R, Jones AL. 2002. Effect of treating apple trees with acibenzolar-*S*-methyl on fire blight and expression of pathogenesis-related protein genes. Plant Disease 86: 785–790.

Meng X, Bonasera JM, Kim JF, Nissinen RM, Beer SV. 2006. Apple proteins that interact with DspA/E, a pathogenicity effector of *Erwinia amylovora*, the fire blight pathogen. Molecular Plant-Microbe Interactions 19: 53–61.

Mine A, Nobori T, Salazar-Rondon MC, Winkelmüller TM, Anver S, Becker D, Tsuda K. 2017. An incoherent feed-forward loop mediates robustness and tunability in a plant immune network. EMBO Reports 18: 464–476.

Monson RK, Trowbridge AM, Lindroth RL, Lerdau MT. 2022. Coordinated resource allocation to plant growth–defense tradeoffs. New Phytologist 233: 1051–1066.

Nieuwenhuizen NJ, Green SA, Chen X, Bailleul EJD, Matich AJ, Wang MY, Atkinson RG. 2013. Functional genomics reveals that a compact terpene synthase gene family can account for terpene volatile production in apple. Plant Physiology 161: 787–804.

Parent B, Turc O, Gibon Y, Stitt M, Tardieu F. 2010. Modelling temperature-compensated physiological rates, based on the co-ordination of responses to temperature of developmental processes. Journal of Experimental Botany 61: 2057–2069.

Paulin J-P, Sampson R. 1973. Le feu bactérien en France. II. Caractères des souches d’*Erwinia amylovora* (Burrill), Winslow et al. 1920, isolées du foyer franco-belge. Annales de Phytopathologie 5: 389–397.

Peng Y, Yang J, Li X, Zhang Y. 2021. Salicylic acid: biosynthesis and signaling. Annual Review of Plant Biology 72: 761–791.

Perrin A, Daccord N, Roquis D, Celton J-M, Vergne E, Bucher E. 2020. Divergent DNA methylation signatures of juvenile seedlings, grafts and adult apple trees. Epigenomes 4: 4.

Pompili V, Dalla Costa L, Piazza S, Pindo M, Malnoy M. 2020. Reduced fire blight susceptibility in apple cultivars using a high-efficiency CRISPR/Cas9-FLP/FRT-based gene editing system. Plant Biotechnology Journal 18: 845–858.

Qiu J, Xie J, Chen Y, Shen Z, Shi H, Naqvi NI, Qian Q, Liang Y, Kou Y. 2022. Warm temperature compromises JA-regulated basal resistance to enhance *Magnaporthe oryzae* infection in rice. Molecular Plant 15: 723–739.

Quint M, Delker C, Franklin KA, Wigge PA, Halliday KJ, Zanten M van. 2016. Molecular and genetic control of plant thermomorphogenesis. Nature Plants 2: 15190.

R Development Core Team. 2025.R: A language and environment for statistical computing.

R Studio Team. 2025.RStudio: Integrated development environment for R.

Ratchaseema MTN, Kladsuwan L, Soulard L, Swangmaneecharern P, Punpee P, Klomsa-ard P, Sriroth K, Keawsompong S. 2021. The role of salicylic acid and benzothiadiazole in decreasing phytoplasma titer of sugarcane white leaf disease. Scientific Reports 11: 15211.

Richard MMS, Knip M, Aalders T, Beijaert MS, Takken FLW. 2020. Unlike many disease resistances, Rx1-mediated immunity to Potato Virus X is not compromised at elevated temperatures. Frontiers in Genetics 11: 417.

Roussin-Léveillée C, Rossi CAM, Castroverde CDM, Moffett P. 2024. The plant disease triangle facing climate change: a molecular perspective. Trends in Plant Science 29: 895–914.

Saeed AI, Sharov V, White J, Li J, Liang W, Bhagabati N, Braisted J, Klapa M, Currier T, Thiagarajan M, et al. 2003. TM4: A free, open-source system for microarray data management and analysis. BioTechniques 34: 374–378.

Sandroni M, Liljeroth E, Mulugeta T, Alexandersson E. 2020. Plant resistance inducers (PRIs): perspectives for future disease management in the field. CABI Reviews 15: 001.

Sasaki-Sekimoto Y, Jikumaru Y, Obayashi T, Saito H, Masuda S, Kamiya Y, Ohta H, Shirasu K. 2013. Basic helix-loop-helix transcription factors JASMONATE-ASSOCIATED MYC2-LIKE1 (JAM1), JAM2, and JAM3 are negative regulators of jasmonate responses in Arabidopsis. Plant Physiology 163: 291–304.

van Schie CCN, Takken FLW. 2014. Susceptibility genes 101: how to be a good host. Annual Review of Phytopathology 52: 551–581.

Schmittgen TD, Livak KJ. 2008. Analyzing real-time PCR data by the comparative *C*T method. Nature Protocols 3: 1101–1108.

Shields A, Yao L, Rossi CAM, Collado Cordon P, Kim JH, AlTemen WMA, Li S, Marchetta EJR, Shivnauth V, Chen T, et al. 2025. Warm temperature suppresses plant systemic acquired resistance by intercepting *N*-hydroxypipecolic acid biosynthesis. The Plant Journal 123: e70374.

Široká J, Brunoni F, Pěnčík A, Mik V, Žukauskaitė A, Strnad M, Novák O, Floková K. 2022. High-throughput interspecies profiling of acidic plant hormones using miniaturised sample processing. Plant Methods 18: 122.

Smyth GK. 2005. limma: Linear models for microarray data. In: Gentleman R, Carey VJ, Huber W, Irizarry RA, Dudoit S, eds. Bioinformatics and Computational Biology Solutions Using R and Bioconductor. New York, USA: Springer, 397–420.

Stevens RB. 1960. Chapter 10 - Cultural practices in disease control. In: Horsfall JG, Dimond AE, eds. Plant Pathology: An Advanced Treatise. New York and London: Academic Press, 357–429.

Subramanian A, Tamayo P, Mootha VK, Mukherjee S, Ebert BL, Gillette MA, Paulovich A, Pomeroy SL, Golub TR, Lander ES, et al. 2005. Gene set enrichment analysis: A knowledge-based approach for interpreting genome-wide expression profiles. Proceedings of the National Academy of Sciences of the United States of America 102: 15545–15550.

Tegtmeier R, Pompili V, Singh J, Micheletti D, Silva KJP, Malnoy M, Khan A. 2020. Candidate gene mapping identifies genomic variations in the fire blight susceptibility genes *HIPM* and *DIPM* across the *Malus* germplasm. Scientific Reports 10: 16317.

Tripathi D, Jiang Y-L, Kumar D. 2010. SABP2, a methyl salicylate esterase is required for the systemic acquired resistance induced by acibenzolar-*S*-methyl in plants. FEBS Letters 584: 3458–3463.

Ullah C, Tsai C-J, Unsicker SB, Xue L, Reichelt M, Gershenzon J, Hammerbacher A. 2019. Salicylic acid activates poplar defense against the biotrophic rust fungus *Melampsora larici-populina* via increased biosynthesis of catechin and proanthocyanidins. New Phytologist 221: 960–975.

Vandesompele J, de Preter K, Pattyn F, Poppe B, van Roy N, de Paepe A, Speleman F. 2002. Accurate normalization of real-time quantitative RT-PCR data by geometric averaging of multiple internal control genes. Genome Biology 3: research0034.1-0034.11.

Vanneste JL (Ed.). 2000. Fire blight: the disease and its causative agent, Erwinia amylovora. Wallingford, UK: CABI.

Vasseur F, Pantin F, Vile D. 2011. Changes in light intensity reveal a major role for carbon balance in *Arabidopsis* responses to high temperature. *Plant*, Cell & Environment 34: 1563–1576.

Velásquez AC, Castroverde CDM, He SY. 2018. Plant–pathogen warfare under changing climate conditions. Current Biology 28: R619–R634.

Walters DR, Ratsep J, Havis ND. 2013. Controlling crop diseases using induced resistance: challenges for the future. Journal of Experimental Botany 64: 1263–1280.

Walters D, Walsh D, Newton A, Lyon G. 2005. Induced resistance for plant disease control: Maximizing the efficacy of resistance elicitors. Phytopathology 95: 1368–1373.

Wang Y, Song S, Zhang W, Deng Q, Feng Y, Tao M, Kang M, Zhang Q, Yang L, Wang X, et al. 2025. Deciphering phenylalanine-derived salicylic acid biosynthesis in plants. Nature 645: 208–217.

Warneys R, Gaucher M, Robert P, Aligon S, Anton S, Aubourg S, Barthes N, Braud F, Cournol R, Gadenne C, et al. 2018. Acibenzolar-*S*-methyl reprograms apple transcriptome toward resistance to rosy apple aphid. Frontiers in Plant Science 9: 1795.

Wickham H. 2016. ggplot2: Elegant Graphics for Data Analysis. New York, USA: Springer-Verlag.

Wu T, Hu E, Xu S, Chen M, Guo P, Dai Z, Feng T, Zhou L, Tang W, Zhan L, et al. 2021.clusterProfiler 4.0: A universal enrichment tool for interpreting omics data. The Innovation 2: 100141.

Zeng Q, Puławska J, Schachterle J. 2021. Early events in fire blight infection and pathogenesis of *Erwinia amylovora*. Journal of Plant Pathology 103: 13–24.

Zhang W, Chen S, Abate Z, Nirmala J, Rouse MN, Dubcovsky J. 2017. Identification and characterization of *Sr13*, a tetraploid wheat gene that confers resistance to the Ug99 stem rust race group. Proceedings of the National Academy of Sciences of the United States of America 114: E9483–E9492.

Zhu Y, Qian W, Hua J. 2010. Temperature modulates plant defense responses through NB-LRR proteins. PLoS Pathogens 6: e1000844.

Zhu B, Zhang Y, Gao R, Wu Z, Zhang W, Zhang C, Zhang P, Ye C, Yao L, Jin Y, et al. 2025. Complete biosynthesis of salicylic acid from phenylalanine in plants. Nature 645: 218–227.

