## Supporting Information for "Welcome pathogens: transient heat dampens immune responses to acibenzolar-*S*-methyl in apple plants"

The following Supporting Information is available for this article:

**Figure S1.** Climatic variables measured at the meteorological station of Beaucouzé (France) for the eleven hottest days recorded between March 2016 and July 2018.

**Figure S2.** Climate dynamics applied in the growth chamber to study the effect of heatwaves on apple response to ASM.

**Figure S3.** Impact of heat and treatment on plant phenotype, and representative photographs of apple seedlings and their typical disease symptoms.

**Figure S4.** Accumulation of high temperatures prior to inoculation impairs the ability of apple grafts of GDDH13 to mount an effective ASM-induced protection against *E. amylovora*.

**Figure S5.** Identification of gene clusters by CAST.

**Figure S6.** Protection provided by ASM to apple seedlings against *E. amylovora* is more strongly dampened by a heatwave posterior to ASM application than by a prior heatwave.

**Figure S7.** Putative apple genes of SA biosynthesis pathway are induced by ASM at high temperature.

**Table S1.** List of genes that are differentially regulated by ASM treatment and transient heat according to the microarray analysis.

**Table S2.** Design of “resistance” and “susceptibility” marker genes.

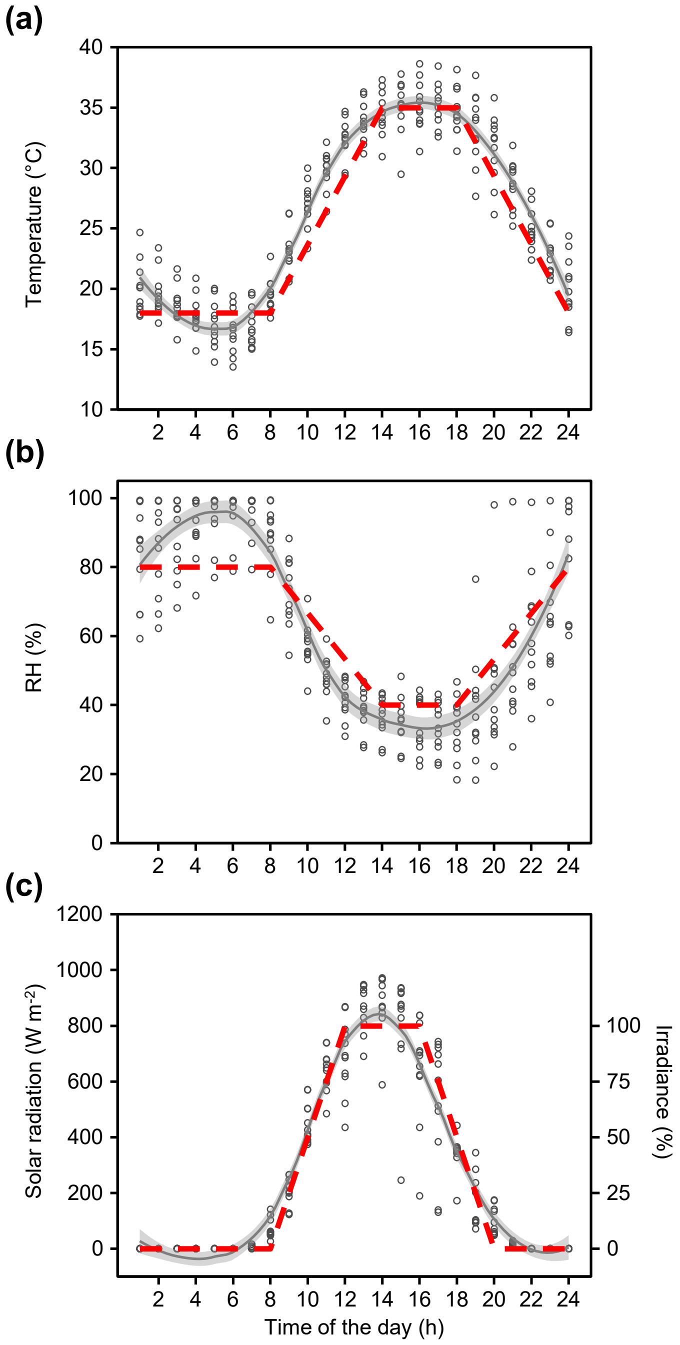


**Fig. S1. Climatic variables measured at the meteorological station of Beaucouzé (France) for the eleven hottest days recorded between March 2016 and July 2018.** Empty circles correspond to individual, hourly values. **(a)** Air temperature, **(b)** relative humidity (RH), **(c)** solar radiation and corresponding percent irradiance. Grey solid curves represent the average trend as determined by the non-parametric loess method (span = 0.5), surrounded by the standard error (grey shaded area). Red dashed lines indicate the piecewise linear approximation of each climatic variable that was used to design the heatwave scenarios in the growth chamber.

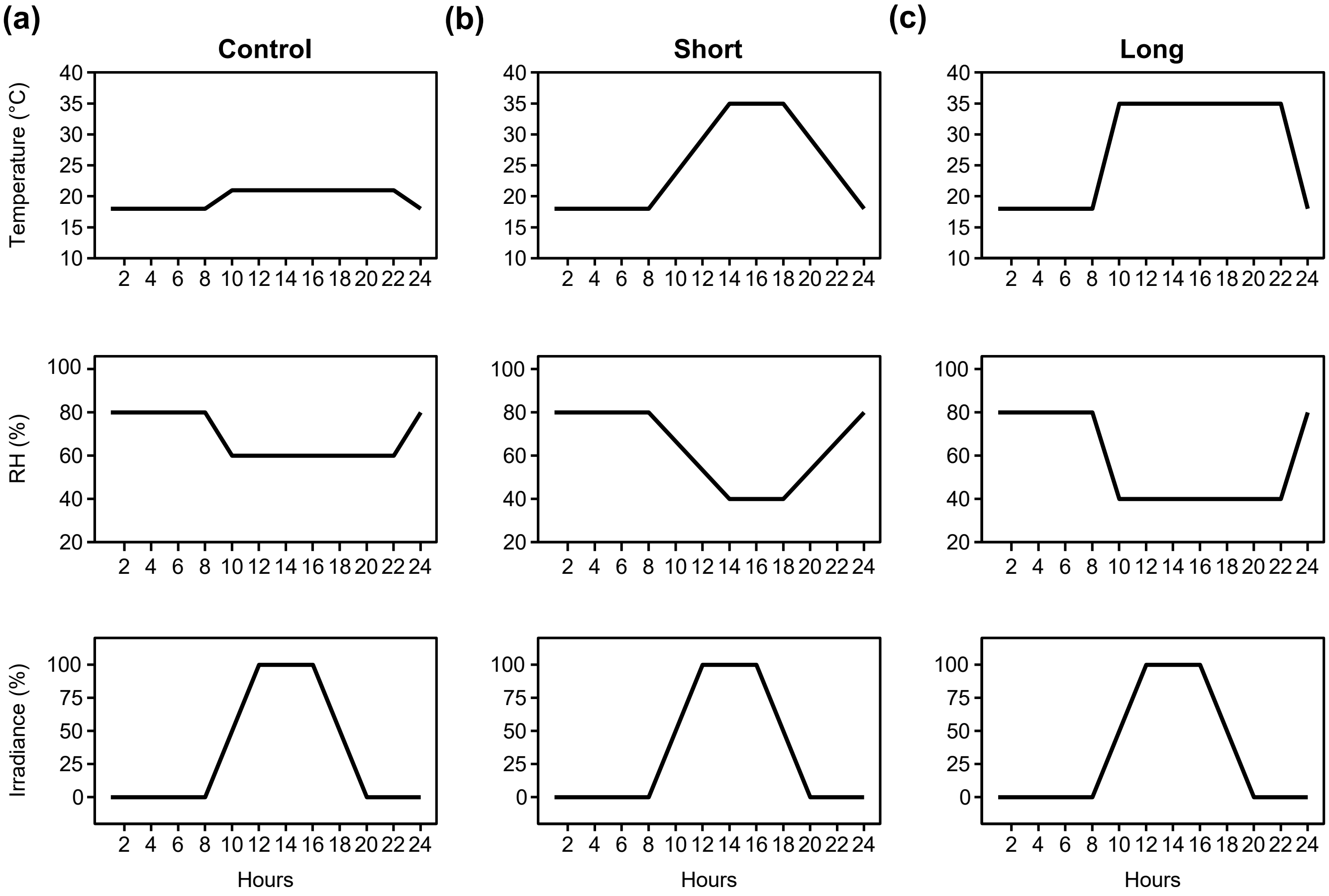


**Fig. S2. Climate dynamics applied in the growth chamber to study the effect of heatwaves on apple response to ASM. (a)** “Control”, **(b)** “Short” and **(c)** “Long” scenarios showing the daily variations in temperature, relative humidity (RH) and percent irradiance applied in growth chambers. Combination of these elemental daily programs were used to reproduce the different thermal scenarios that were studied.

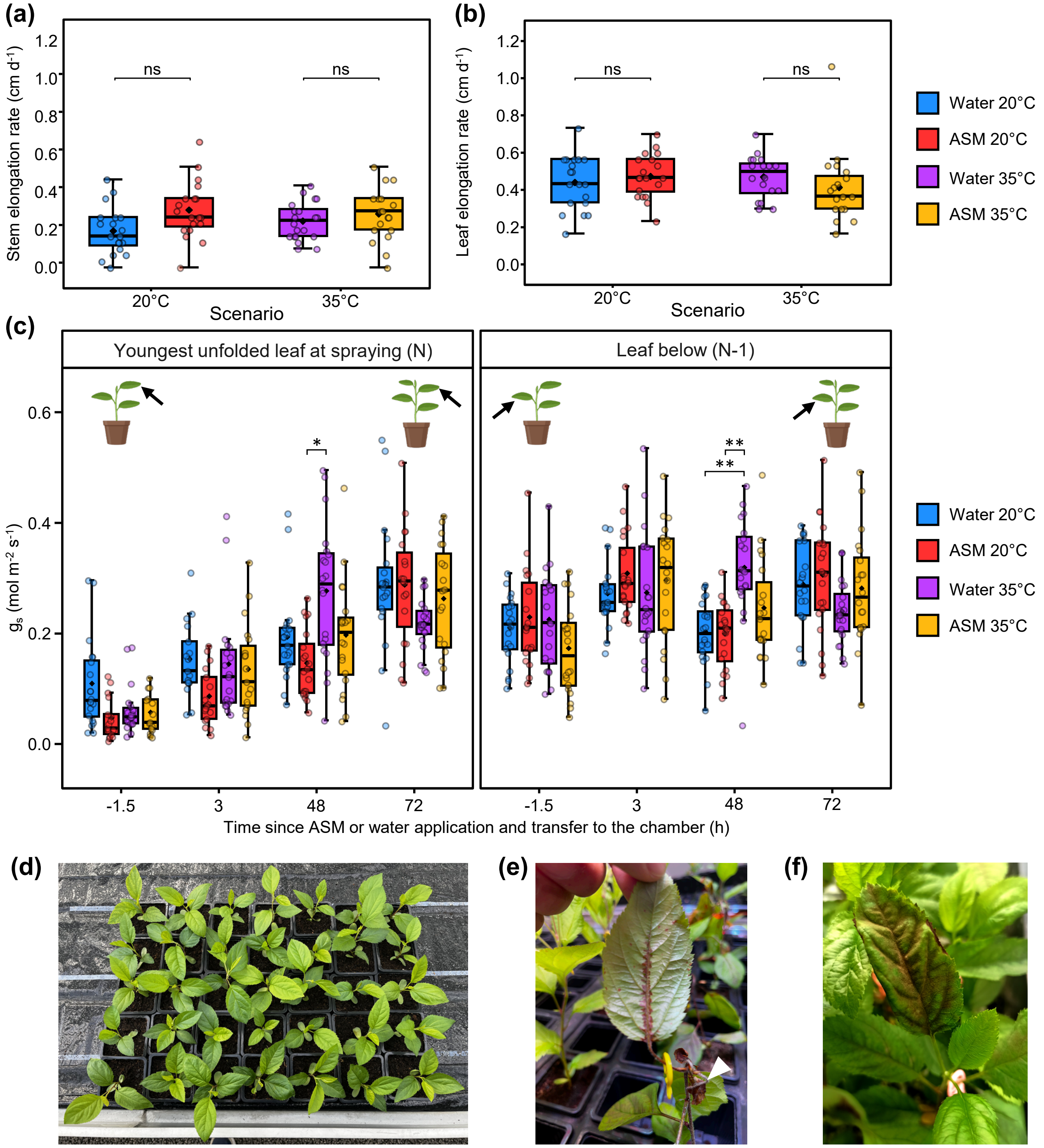


**Fig. S3. Impact of heat and treatment on plant phenotype, and representative photographs of apple seedlings and their typical disease symptoms. (a)**, **(b)**, and **(c)** Boxplots and jitter plots showing the impact of temperature scenario and ASM or water treatment on apple seedling phenotypes. All measurements were performed on the same plants of the same experiment (details in Methods). At time 0, plants were sprayed with ASM or water and immediately transferred from a greenhouse to a growth chamber with controlled environmental conditions. Air temperature was maintained at either 20°C or 35°C for 24 h, and all plants then remained in the growth chamber at 20°C during 48 h more. The elongation rate of **(a)** the stem and **(b)** the youngest unfolded leaf was calculated using measurements made before treatment and 3 d after treatment. Statistical differences between ASM and water treatment were determined at each temperature using pairwise t-tests adjusted by the Bonferroni method, and were deemed not significant (ns). **(c)** Stomatal conductance ($g_s$) was measured on the youngest unfolded leaf at time of ASM/water treatment application (rank N, left panel) and on the leaf rank immediately below (rank N-1, right panel), 1.5 h before treatment and 3, 48 and 72 h after treatment. Multiple mean comparisons were performed between each combination of both factors at each time point using pairwise t-tests adjusted by the Bonferroni method (* $0.05 \geq P > 0.01$, ** $0.01 \geq P > 0.001$). Note that non-significant differences have been omitted for clarity. **(d)** Photograph of a representative batch of apple seedlings at the start of the experiment. **(e)** Photograph of systemic fire blight necrosis symptoms on a diseased apple seedling, 21 d after inoculation by *E. amylovora*. The inoculated leaf is indicated by a white arrowhead and the picture focuses on necrotic symptoms spreading along the midvein of a distant leaf, and along the stem up to the plant apex. **(f)** Photograph of an apple seedling leaf showing *V. inaequalis* sporulation, 21 d after inoculation.

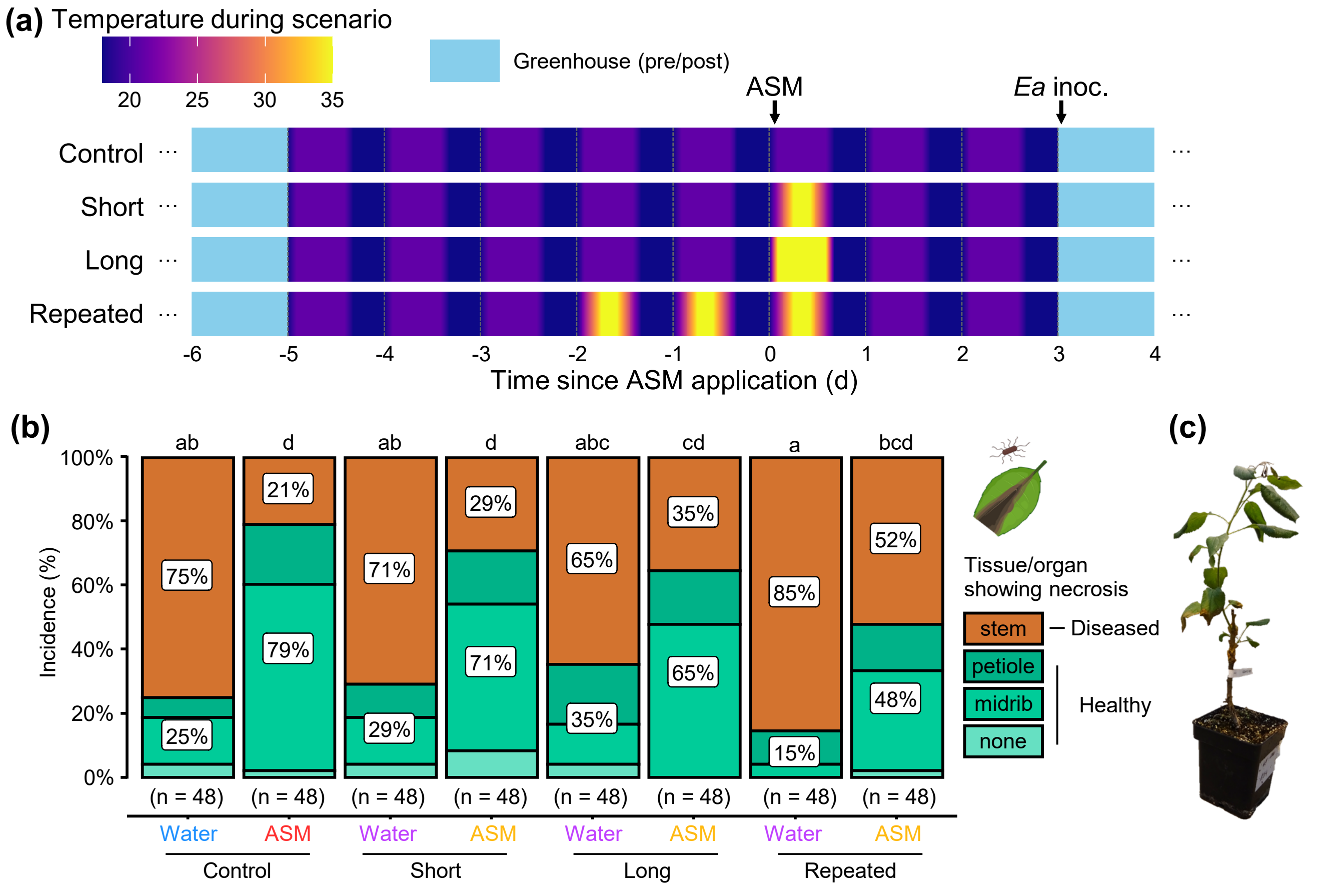


**Fig. S4. Accumulation of high temperatures prior to inoculation impairs the ability of apple grafts of GDDH13 to mount an effective ASM-induced protection against *E. amylovora*. (a)** Schematic representation of the experimental protocol (details in Methods). Air temperature during the contrasting thermal scenario (period in the growth chamber) is color-coded according to the color map. Arrows indicate the application of ASM (or water) treatments and pathogen inoculation on the youngest unfolded leaf. **(b)** Stacked bar chart showing the proportion of healthy (asymptomatic on stem) and diseased (symptomatic on stem) grafted plants inoculated with *E. amylovora*, 28 d post-inoculation. Data represent 48 biological replicates from 4 independent trials (n = total number of plants). Statistical significance was determined using pairwise Fisher's exact tests; same letters indicate groups that are not significantly different. Icons created in BioRender.com. **(c)** Photograph of a diseased GDDH13 graft presenting fire blight symptoms on the apex and displaying the typical “hook shape”.

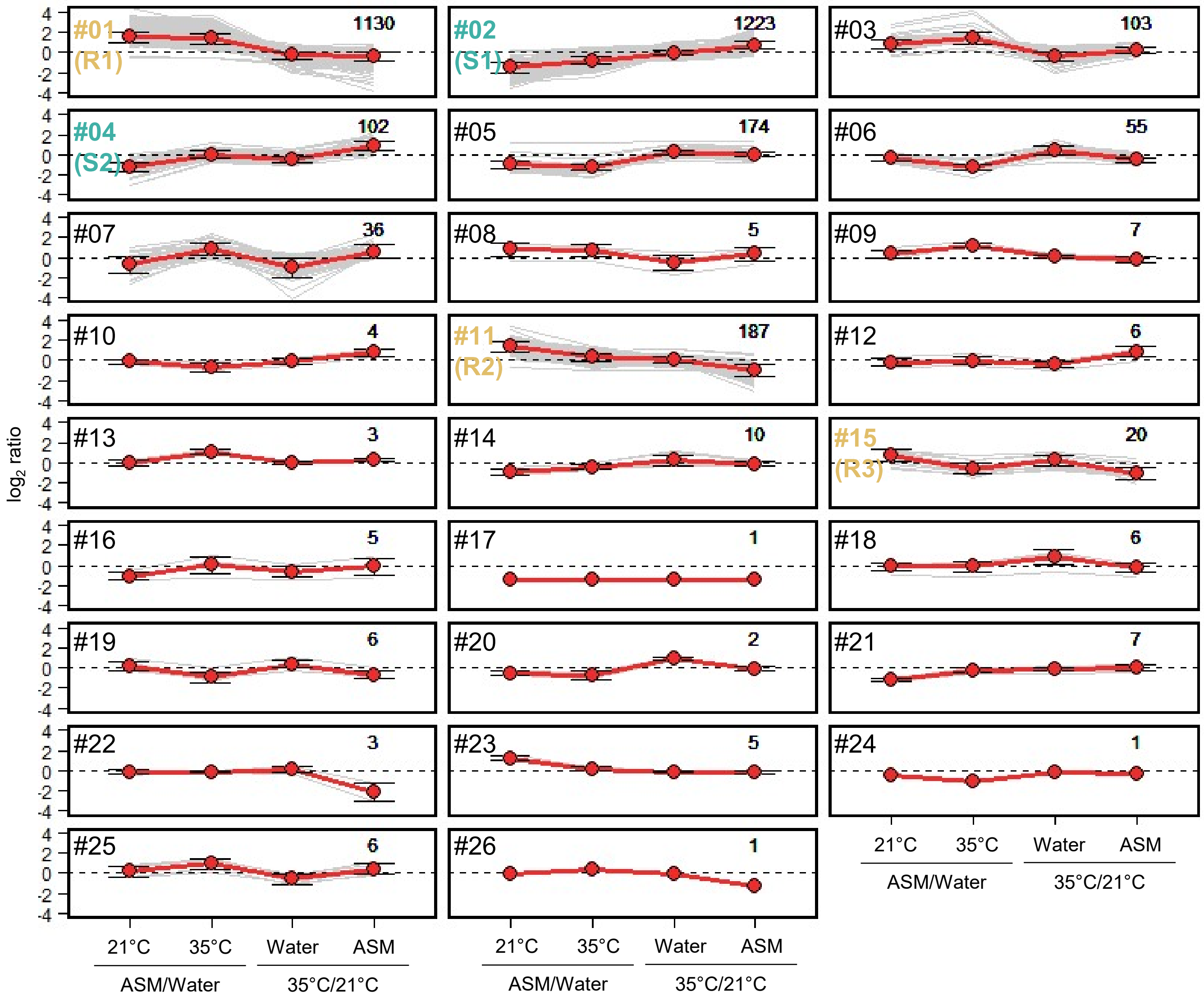


**Fig. S5. Identification of gene clusters by CAST.** Clustering was based on the expression pattern of 3108 genes that displayed differential expression (defined by a $|\log_2 \text{ratio}| \geq 1$ and $P < 0.05$ in at least one microarray comparison, see Fig. 5b) in plants treated with ASM or water and exposed to control (21°C) or elevated (35°C) temperature. Pearson correlation distance metric (0.95 threshold affinity) allowed the construction of 26 clusters. Grey lines correspond to the expression ratio pattern of each individual DEG. Red points correspond to the average $\log_2$ expression ratio (± sd) within each comparison, for each cluster. The number of DEGs grouped in each cluster and the corresponding cluster number (and name for R1, R2, R3, S1 and S2 investigated in Fig. 6) are indicated in the upper right- and left-hand corner of each box, respectively.

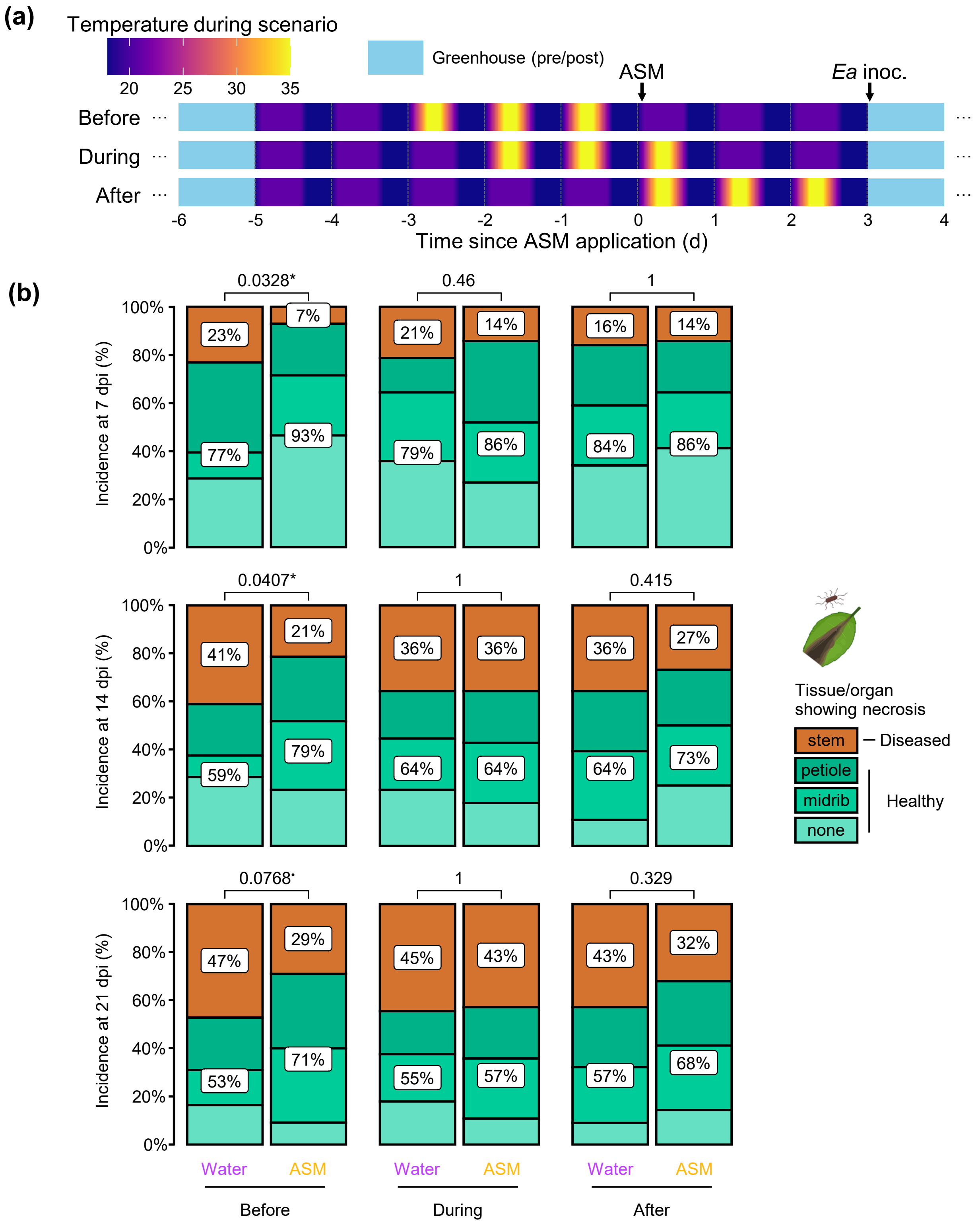


**Fig. S6. Protection provided by ASM to apple seedlings against *E. amylovora* is more strongly dampened by a heatwave posterior to ASM application than by a prior heatwave. (a)** Schematic representation of the experimental protocol (details in Methods). Air temperature during the contrasting thermal scenario (period in the growth chamber) is color-coded according to the color map. Arrows indicate the application of ASM (or water) treatments and pathogen inoculation on the youngest unfolded leaf. **(b)** Stacked bar charts showing the proportion of healthy (asymptomatic on stem) and diseased (symptomatic on stem) seedlings inoculated with *E. amylovora* at 7, 14 and 21 d post-inoculation (dpi). Data represent 55–56 biological replicates (number of plants in each condition) from a single trial. The *p*-value of Fisher's exact test between water and ASM treatments is shown above each pair of barplot (•P<0.1, *P<0.05). Icons created in BioRender.com.

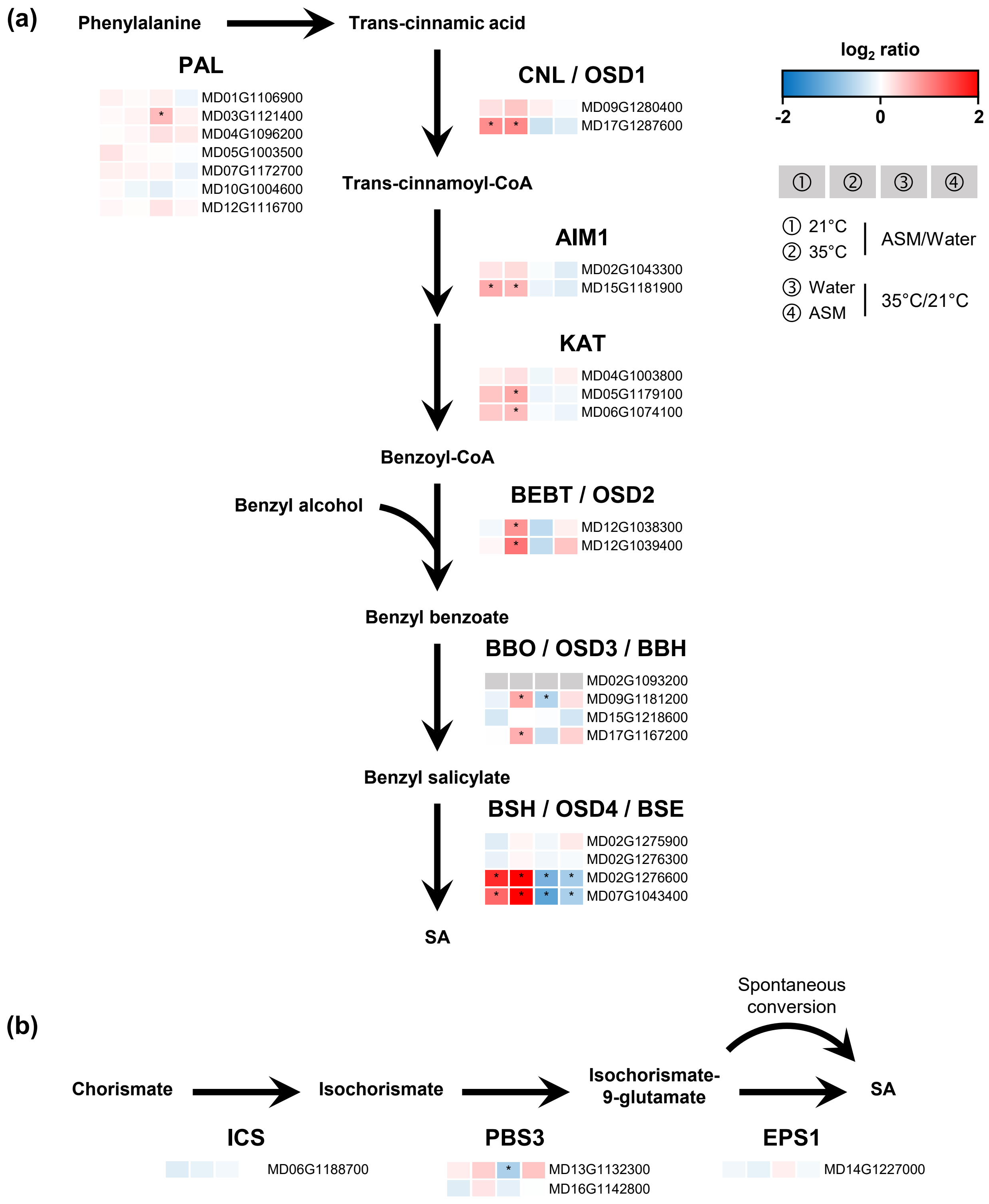


**Fig. S7. Putative apple genes of SA biosynthesis pathway are induced by ASM at high temperature.** The expression pattern of putative apple orthologues of genes involved in SA biosynthesis was extracted from microarray data on GDDH13 leaves (Fig. 5). For each enzymatic step, expression of corresponding gene(s) are color-coded in boxes showing the effect of ASM ① at 21°C and ② at 35°C, and the effect of high temperature upon treatment with ③ water and ④ ASM. Significant up- or down-regulations are notified (*). The gene represented by grey boxes was absent from the microarray. **(a)** Phenylalanine ammonia-lyase (PAL) pathway. Apple protein orthologues of cinnamoyl-coenzyme A ligase (CNL) / *Oryza sativa* SA-DEFICIENT GENE 1 (OSD1), ABNORMAL INFLORESCENCE MERISTEM1 (AIM1), 3-ketoacyl-CoA thiolase (KAT), benzyl alcohol benzoyltransferase (BEBT) / OSD2, benzyl benzoate oxidase (BBO) / ODS3 / benzylbenzoate hydroxylase (BBH) and benzyl salicylate hydrolase (BSH) / OSD4 / benzylsalicylate esterase (BSE) were identified according to a BLAST of protein sequence from rice. Apple protein with an e-value of 0, less than 10% of gaps and 60% of identities were retained. If no protein met these criteria, the top four hits were selected. **(b)** Isochorismate synthase (ICS) pathway. Apple mRNA orthologues of ICS, AVRPPHB SUSCEPTIBLE 3 (PBS3) and ENHANCED PSEUDOMONAS SUSCEPTIBILITY 1 (EPS1) were identified according to a BLAST with at least 60% of identities with CDS sequence from *Arabidopsis thaliana*.

**Table S1. List of genes that are differentially regulated by ASM treatment and transient heat according to the microarray analysis.** Differentially expressed genes were defined by a $|\log_2 \text{ratio}| \geq 1$ and $P < 0.05$. The table shows gene accession, gene description, gene symbol (if any), $\log_2$ fold change within each comparison, cluster membership (see Fig. S5), and functional annotation. Gene description and functional annotation are based on the GDDH13 genome annotation (Daccord et al., 2017). Available online as an Excel file.

**Table S2. Design of "resistance" and "susceptibility" marker genes.** Genes initially selected through CAST belonged to "resistance-associated" (R1, R2, R3) or "susceptibility-associated" (S1, S2) clusters (Fig. 6). Homologous genes were searched in the genome, and were eventually found in the same or an alternative cluster. Primers were designed to target as many homologous genes as possible, and the targeted ones are bolded. The table shows gene name, functional annotation, accession, cluster membership (if any), and primer sequence and concentration. Functional annotation is based on the GDDH13 genome annotation (Daccord et al., 2017). Available online as an Excel file.
